# Chromokinesin Klp-19 regulates microtubule overlap and dynamics during anaphase in *C. elegans*

**DOI:** 10.1101/2023.10.26.564275

**Authors:** Vitaly Zimyanin, Magdalena Magaj, Nadia Ingabire Manzi, Che-Hang Yu, Theresa Gibney, Yu-Zen Chen, Mustafa Basaran, Xavier Horton, Karsten Siller, Ariel Pani, Daniel Needleman, Daniel J. Dickinson, Stefanie Redemann

## Abstract

Recent studies have highlighted the significance of the spindle midzone, the region between the segregating chromosomes, in ensuring proper chromosome segregation. By combining 3D electron tomography, cutting-edge light microscopy and a novel single cell *in vitro* essay allowing single molecule tracking, we have discovered a previously unknown role of the regulation of microtubule dynamics within the spindle midzone of *C. elegans* by the chromokinesin KLP-19, and its relevance for proper spindle function. Using Fluorescence recovery after photobleaching and a combination of second harmonic generation and two-photon fluorescence microscopy, we found that the length of the antiparallel microtubule overlap zone in the spindle midzone is constant throughout anaphase, and independent of cortical pulling forces as well as the presence of the microtubule bundling protein SPD-1. Further investigations of SPD-1 and KLP-19 in *C. elegans*, the homologs of PRC1 and KIF4a, suggest that KLP-19 regulates the overlap length and functions independently of SPD-1. Our data shows that KLP-19 plays an active role in regulating the length of microtubules within the midzone as well as the size of the antiparallel overlap region throughout mitosis. Depletion of KLP-19 in mitosis leads to an increase in microtubule length and thus microtubule-based interactions in the spindle midzone, which affects spindle dynamics and force transmission. Our data shows that by localizing KLP-19 to the spindle midzone in anaphase microtubule dynamics can be locally controlled allowing the formation of a functional midzone.

**Summary:** KLP-19 controls microtubule length in the spindle midzone of *C. elegans*, affecting spindle dynamics and force transmission during mitosis.

## Introduction

Anaphase, the process that separates the duplicated genetic material, consists of two different phases: Anaphase A, a shortening of the chromosome to spindle pole distance, and Anaphase B, an increase in the distance between the spindle poles. The forces that drive chromosome segregation during anaphase A are thought to be generated by depolymerization of kinetochore microtubules, while the forces in anaphase B are thought to rely on a combination of antiparallel sliding of microtubules (MT) between the chromosomes and cortical pulling forces on astral MTs, both driven by motor proteins. In many spindles, both anaphase A and anaphase B occur simultaneously (*1*).

Chromosome segregation in the first mitotic division of the *C. elegans* embryo has long been known to be strongly affected by cortical force generators, a trimeric complex composed of a membrane anchored Gα-protein GOA-1/GPA-16, GPR-1/2 and LIN-5 that recruits dynein, during anaphase. Those force generators generate pulling forces that elongate the spindle (*2–5*) and which were thought to drive chromosome segregation. As the pole-to-chromosome distance remains constant during this process, Anaphase B was proposed to be the main mechanisms in *C. elegans* (*6*). However, it was recently shown that spindle elongation and chromosome segregation are two mechanistically distinct processes in *C. elegans* mitosis: spindle elongation results from cortical pulling forces (*2–5, 7*), while chromosome segregation is presumably primarily governed by poorly appreciated processes internal to the spindle (*7–9*). Analysis of the midzone in tissue culture cells has confirmed a similar role for the spindle midzone during chromosome segregation in mammalian cells (*7*). However, we still lack significant information about how the spindle midzone generates forces contributing to chromosome segregation during anaphase (*10*).

Several conserved proteins localize to the spindle midzone in mammalian cells and *C. elegans,* coordinating its assembly and function. BUB1/BUB-1, a kinetochore protein kinase, was suggested to control the initiation of midzone assembly together with centromere protein CENPF/HCP-1/2 and CLASP/CLS-2. Several publications indicated that the microtubule regulator CLS-2 promotes the *de novo* nucleation of microtubules in the midzone, thus possibly driving the formation of the spindle midzone in *C. elegans* (*9, 11–13*). In addition, the RAN pathway has been suggested to play a role in the midzone by promoting microtubule assembly or stabilization (*14–19*). The microtubule bundling factors PRC1/ SPD-1 the kinesins KIF4a/ KLP-19 and EG5/ BMK-1 as well as centralspindlin, a protein complex composed of the kinesin MKLP1/ ZEN-4 and RACGAP1/ CYK-4, then localize to the midzone where they regulate the microtubule organization (*20, 21*).

A crucial component of midzone stability and function in mammalian cells is PRC1, which plays a central role in crosslinking antiparallel microtubules (*22*, *23–26*). The absence of PRC1 leads to an inability to establish robust spindle midzones and frequent cytokinesis failures (*23, 27–29*).

Depletion of SPD-1, the PRC1 homolog in *C. elegans* similarly prevents the formation of a stable midzone, leading to premature spindle rupture. Cytokinesis however is mostly completed, presumably due to contractile ring constriction-driven bundling of astral microtubules at the furrow tip that leads to the formation of a midbody in the absence of a spindle midzone (*30*).

PRC1 has been shown to directly interact with various components of the midzone, such as CLASP1, a protein that binds to the plus-ends of microtubules and promotes their growth (*13, 31–33*), MKLP1, a subunit of the centralspindlin complex, contributing to the mechanical properties of the midzone (*34*, *35*), and the kinesin KIF4A, which inhibits the elongation of microtubule plus-ends within the midzone (26, *36–38*). In *C. elegans* SPD-1 was also shown to directly interact with CYK-4, a component of the centralspindlin complex (*34*), but other interactions in *C. elegans* have not yet been assessed.

*In vitro* experiments on individual microtubules showed that PRC1 and KIF4A accumulate to form dynamic end tags, with the length of these tags being proportional to the concentration of PRC1 and the length of the microtubules (*38, 39*). Mixtures of PRC1, KIF4A, and microtubules also give rise to antiparallel bundles in vitro (*40–42*). As microtubules slide apart, the overlap zones between them shorten and eventually reach a steady state, resembling the length of overlap zones within the midzone observed in cells (*40, 41*). PRC1 has been shown to generate frictional forces that counteract microtubule sliding, aligning with the braking function of midzones observed in cells (*43–45*).

In *C. elegans*, depletion of either SPD-1/PRC1, BMK-1/ EG5 or ZEN-4/ MKLP1 results in rapid chromosome segregation and often breakage of the spindle midzone during anaphase. The spindle rupture can be suppressed by depletion of cortical pulling forces, suggesting that those midzone proteins are required for the mechanical integrity of the midzone during its elongation (*9, 13, 34*). These observations strongly indicate a dual role for the midzone in anaphase. Firstly, it acts as a stabilizing element, countering intense cortical pulling forces and regulating the rate of chromosome segregation. Secondly, it functions as a force generator, facilitating chromosome segregation (*7, 8*). This complex interplay underscores the challenge of balancing the conflicting demands on the midzone during cell division. The mechanisms how the spindle midzone achieves this have remained elusive.

Combining light and electron microscopy, as well as a novel *in vitro* microtubule assay utilizing cell lysate of single, staged *C. elegans* embryos expressing fluorescently tagged KLP-19 and SPD-1 we show that the chromokinesin KLP-19 is a processive motor, which regulates the dynamics, stability and length of microtubules in the spindle midzone during anaphase. Our data suggests that KLP-19 regulates the microtubule overlap in the spindle midzone independently of SPD-1 and that both proteins do not directly interact. In absence of KLP-19 microtubules are longer, leading to an increase in microtubule overlap in the spindle midzone and thus enabling more interactions with neighboring microtubules. Our data suggests that the increase in the microtubule overlap affects the force transmission in the spindle, leading to a decrease in pole and chromosome separation. In summary, by localizing KLP-19 to the spindle midzone in anaphase microtubule dynamics can be locally controlled allowing the formation of a functional midzone.

## Results

### Microtubule overlap in the midzone remains constant throughout anaphase

An active role of the spindle midzone in promoting and restricting chromosome segregation requires a very tight regulation of its structure and dynamics. We previously used 3D electron tomography to reconstruct the spindle midzone in anaphase (*7*) these reconstructions showed that some microtubules in the midzone seem to originate from the region between chromosomes and poles, while other microtubules have both ends between the segregating chromosomes, occasionally contacting the inner-surface of the chromosomes, and potentially nucleating between the chromosomes. Interestingly, only 1 or 2 microtubules extended all the way from pole to pole.

Outward sliding of antiparallel midzone microtubules is thought to be the main regulator of midzone driven chromosome segregation. To quantify changes in the antiparallel overlap of microtubules in the spindle midzone throughout anaphase in the one-cell stage *C. elegans* embryo we used second harmonic generation (SHG) microscopy and two photon florescence (TP) microscopy simultaneously to measure the polarity of the collective microtubules throughout the spindle in vivo (*46*). SHG is a nonlinear optical process in which highly polarizable, non-centrosymmetric materials emit photons with half the wavelength of incident light. When an array of microtubules is imaged with SHG microscopy, the resulting SH signals depend on both the polarity of the microtubule array and the density of microtubules within a focal volume. The readout of microtubule density can be obtained using TP microscopy and was used to compute microtubule polarity from the SHG signals (*46*). The polarity of microtubules ranges from 0 to 1 continuously: 1 corresponds to parallel microtubules with all the plus ends pointing in the same direction, and 0 corresponds to antiparallel microtubules with equal number of plus ends pointing in the opposite direction. Measurements by SHG and TP in wild type embryos showed that microtubules have a constant antiparallel overlap length in the central spindle (Fig. 1A), indicated by the constant width of the polarity curve during anaphase. To test the potential impact of cortical pulling forces on the size of the antiparallel microtubule region in the midzone we repeated the experiment in embryos depleted of GPR-1/2. The size of the antiparallel region did not change in the absence of pulling forces in comparison to control and over time (Fig. 1B), showing that the antiparallel overlap region is constant throughout anaphase and independent of pulling forces. These results suggest that in addition to microtubule sliding, some active process maintains the extent of microtubule overlap during mitosis.

**Figure 1.**
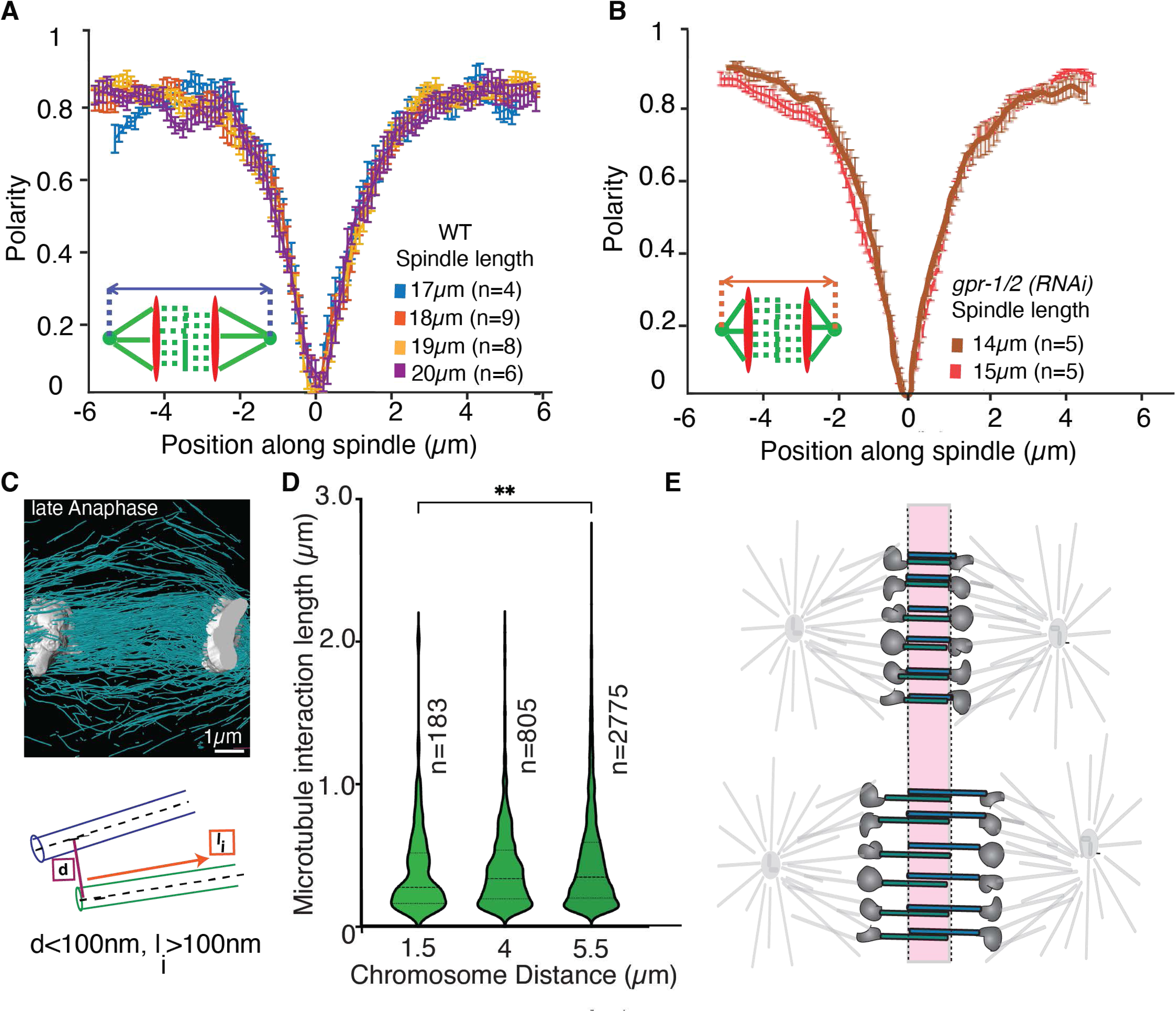
The microtubule overlap in the midzone is constant throughout anaphase. **A.** Polarity plot of microtubules in the spindle at different timepoints in anaphase obtained by the combination of SHG and TP microscopy. 0 on the x-axis represents the spindle center. Spindle length is indicated in the legend as a measure of progression throughout anaphase. **B.** Polarity plot of microtubules in the spindle throughout anaphase after *gpr-1/2 (RNAi)* obtained by SHG and TP microscopy combined. **C.** Top: Representative image of a 3D tomographic reconstruction of the spindle midzone in anaphase in the *C. elegans* embryo, focusing on the region between the chromosomes. Microtubules are shown in cyan, chromosomes in grey. Bottom: Cartoon of a microtubule-microtubule interaction showing the different parameters used to define interactions. d = center to center distance between two microtubules, l_i_ = length of interaction. **D.** Violin Plot of the average interaction length l_i_ between microtubules with maximum d= 100nm and minimum l_i_ = 100nm for 3 different datasets throughout anaphase, chromosome distance as an indicator for progression through anaphase is provided. Number of detected interactions is indicated as n. **E.** Cartoon summarizing the data, showing that microtubule overlap remains constant throughout anaphase in *C. elegans*.

In addition to the SHG analysis we also quantified the length of interactions between individual microtubules in 3 tomographic reconstructions of mitotic spindles, two of which were previously published (*7*), at different stages in anaphase defined by the distance between the segregating chromosomes (Fig. 1C). The interactions were defined by a maximum center-to-center distance of 100 nm between interacting microtubules, based on our previous data on the spindle network (*47*) as well as the reported sizes of microtubule-associated proteins or molecular motors (*48–50*) (Fig. 1C). We further required that this distance was maintained over a minimum length of 100 nm (based on the detection limit of microtubules, which is 100nm). We did not restrict the interaction angle; it is however important to point out that nearly all of the microtubules in the midzone are parallel to each other. Our analysis of microtubule overlap in the midzone (Fig. 1D) showed that the average length of the interactions between midzone microtubules is 390 nm ± 440 nm (STD) in early anaphase (chromosome distance 1.5µm), 408 nm ± 286 nm (STD) in mid anaphase (distance 4µm) and 463 nm ± 368 nm (STD) in late anaphase (distance 5.5µm). While the average length increased significantly between the early and late anaphase stage (based on chromosome distance), the interaction length of microtubules increased by approximately 70 nm between MTs from early to late anaphase, while the size of the central spindle, defined by the distance between the segregating chromosomes, increased by ∼4μm (60x more) at the same time (Fig. 1D). These data suggest that the microtubule overlap in the *C. elegans* midzone does not undergo major changes in length throughout anaphase (Fig. 1E), in contrast to mammalian cells (*48, 51–56*).

### Depletion of SPD-1 and KLP-19 affect spindle dynamics and chromosome segregation in *C. elegans*

The interaction of the kinesin KIF4a with the microtubule bundling protein PRC1 has been suggested to be key to the regulation of microtubule overlap. Based on *in vitro* data it has been proposed that PRC1 recruits KIF4a to the microtubules in the spindle midzone, where it regulates microtubule dynamics and midzone length, possibly through its ability to inhibit microtubule plus-end growth (*22, 26*).

During *C. elegans* mitosis the PRC1 homolog, SPD-1, localizes to the spindle midzone in anaphase (*57*). Our analysis of spindle elongation (pole to pole distance) (Fig. 2A,B, Movie 1, 2) and chromosome segregation (chromosome distance) (Fig. 2A,C) showed that metaphase spindles were shorter after *spd-1 (RNAi)* (*control:* 15.52µm ± 0.26 µm, (n=26), SPD-1 (13.26µm ± 0.35µm, (n=20), sem) as well as SPD-1/ GPR-1/ 2 co-depletion (12.03µm ± 0.25µm; (n=9)) (Fig. 2A,B, Suppl. Fig. 1, Movie 3). Depletion of SPD-1 alone lead to increased rates of spindle elongation and chromosome segregation due to spindle rupture during anaphase (Fig. 2 D,E, Suppl. Fig. 1) (control: elongation 0.09µm/s ± 0.004, segregation 0.13 µm/s ±0.002, *spd-1 (RNAi)* elongation 0.30µm/s ± 0.04, segregation 0.23 µm/s ±0.), as previously reported (*57*). Co-depletion of SPD-1 and GPR-1/ 2 prevented spindle rupture and the rate of spindle elongation was significantly decreased (0.059µm/s± 0.003) in comparison to control embryos as well as *spd-1 (RNAi)* alone (Fig. 2A-E, Suppl. Fig. 1). The rate of chromosome segregation in *spd1/ gpr-1/ 2 (RNAi)* embryos (0.11µm/s ±0.004) was also reduced (Fig. 2E).

**Figure 2.**
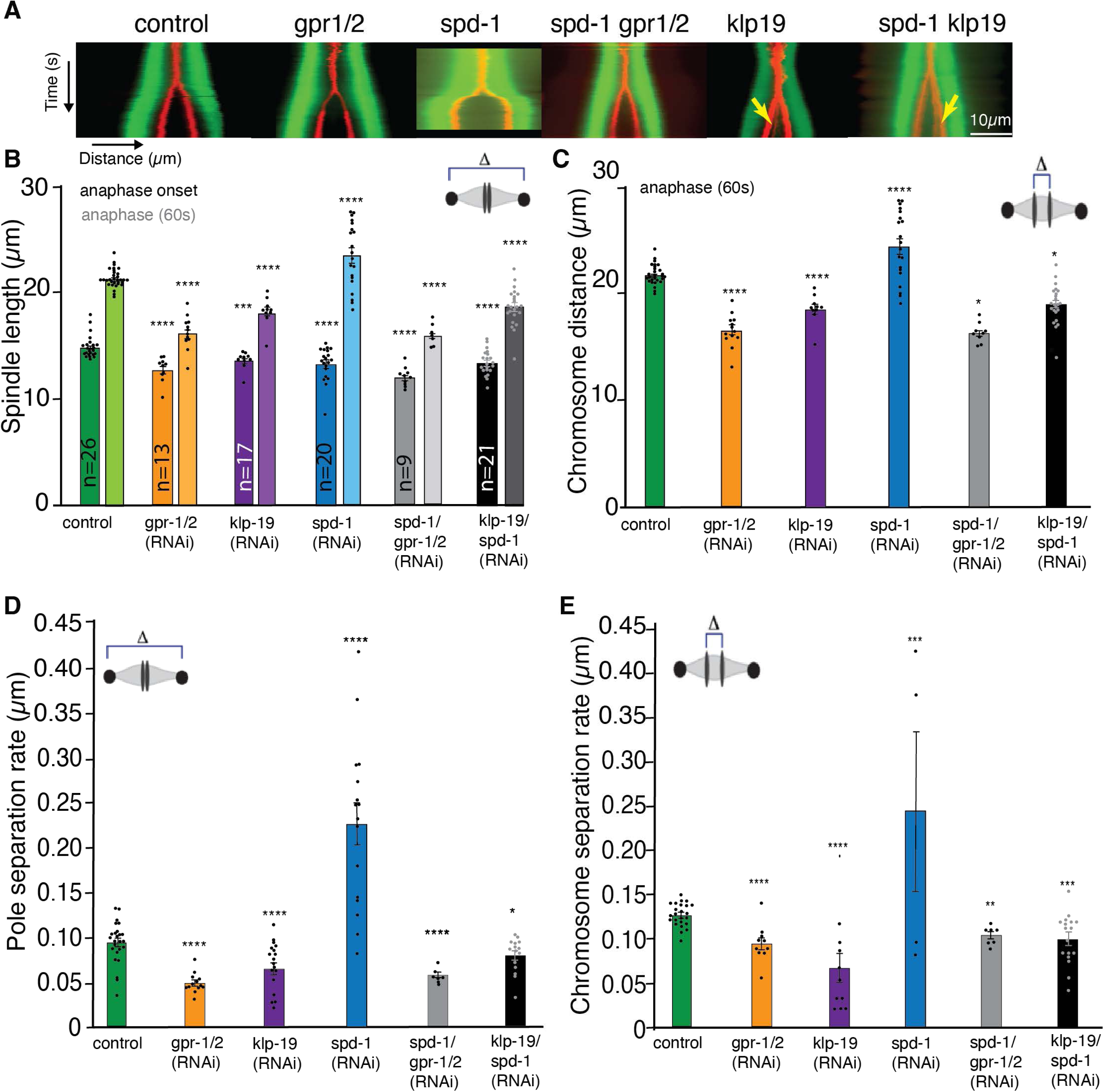
Depletion of KLP-19 and SPD-1 affect spindle dynamics. **A.** Kymographs of *C. elegans* one-cell embryos from metaphase to anaphase in control and after *gpr-1/2, spd-1, spd-1/gpr-1/2, klp-19, and spd-1/klp-19 (RNAi)* obtained by light microscopy. Microtubules are labeled in green, Chromosomes in red. Yellow errors point to lagging chromosomes. Scale Bar 10µm. **B.** Bar plot of the spindle length at anaphase onset (last frame before visible chromosome segregation) and 60s after anaphase onset in control embryos and after different RNAi treatments. **C**. Bar plot of the Chromosome distance 60s after anaphase onset in control embryos and after different RNAi treatments. **D.** Bar Plot of pole-to-pole segregation rate throughout anaphase in control embryos and embryos after different RNAi treatments **E.** Bar Plot of chromosome segregation rate throughout anaphase in control embryos and embryos after different RNAi treatments. Color code: control embryos = green, *gpr-1/2 (RNAi)* = orange, *klp-19 (RNAi)* = purple, *spd-1 (RNAi)* = blue, *spd-1/ gpr-1/2 (RNAi)* = grey, *klp-19/ spd-1 (RNAi)* = black. Error Bars are sem. The significance of differences between control and RNAi conditions was determined by two-tailed Student’s *t*-tests (\*\*\*\**P* < 0.0001).

Depletion of the KIF4a *C. elegans* homolog KLP-19 resulted in severe chromosome segregation defects and showed a strong disruption in metaphase plate and kinetochore alignment and its overall organization, including reduced compaction of chromosomes, as well as presence of multiple lagging chromosomes during segregation (Suppl. Fig 2). We confirmed a role for KLP-19 during mitosis, in generating polar-ejection forces that move chromosomes away from the spindle pole towards the metaphase plate during congression (Suppl Fig 3) (*58*). The observed chromosome missegregation had been associated with errors in the correct formation of end-on connections of MTs to kinetochores (*58*). Detailed analysis of the chromosome segregation and spindle elongation rates showed that *klp-19 (RNAi)* leads to the formation of shorter metaphase spindles (13.71µm ± 0.18µm, (n=17), sem) (Fig 2A,B, Suppl. Fig. 1, Movie 4), and a reduction in the pole-to-pole and chromosome segregation rates (elongation 0.07µm/s ± 0.006 µm/s and segregation 0.08µm/s± 0.008) (Fig 2D, E). To further analyze the role of KLP-19 and SPD-1 we co-depleted both proteins by RNAi. Chromosome congression errors and subsequent chromosome missegregation still occurred upon co-depletion (Suppl Fig 1, Suppl. Fig 2).

However, co-depletion of SPD-1 and KLP-19 did not lead to spindle rupture as observed by *spd-1 (RNAi)* alone (Fig. 2A-E, Suppl.Fig. 1, Suppl. Fig. 2, Movie 5). In contrast, spindle rupture was prevented and spindle elongation rates, as well as chromosome segregation rates were slightly reduced in comparison to control embryos (*elongation* 0.08µm/s ± 0.004, segregation 0.10µm/s± 0.007, (n=21)) (Fig 2A-E). As spindle rupture in SPD-1 depleted embryos is dependent on cortical pulling forces, we quantified the presence of pulling forces after *klp-19 (RNAi)* to assess potential effects on those forces related to the absence of KLP-19. For this we used laser ablation to sever the posterior centrosome from the spindle body and subsequently tracked the velocity of the anterior and posterior centrosomes. This data showed that pulling forces are still present in embryos depleted of KLP-19 (Suppl. Fig. 4) and are not the underlying cause of the lack of spindle rupture upon SPD-1 and KLP-19 double depletion. Thus, KLP-19 and cortical force generators play parallel roles in generating forces that are resisted by SPD-1.

Another possibility is that the lagging chromosomes after *klp-19 (RNAi)* could be responsible for the decrease in spindle length and chromosome segregation as well as preventing spindle rupture after SPD-1 depletion. To test this, we compared the spindle dynamics of *klp-19 (RNAi)*, *spd-1 (RNAi)*, and *spd-1/ klp-19 (RNAi)* treated embryos with embryos depleted of HCP-6. HCP-6 is part of the SMC (Structural Maintenance of Chromosomes) complex in *C. elegans* and plays a crucial role in chromosome segregation during cell division (*59–61*). Specifically, it is involved in maintaining chromosome rigidity and proper orientation during mitosis (*59, 61*). Depleting HCP-6 in *C. elegans* leads to improper chromosome alignment, merotelic kinetochore microtubule attachments and segregation errors, causing aneuploidy. Quantification of spindle length throughout mitosis revealed that spindles in all tested conditions were of similar length at the timepoint of Nuclear envelope breakdown (NEBD) (control: 11.6µm ± 0.3µm, (n=48); *klp-19 (RNAi)* 11.3µm ± 0.3µm, (n=19); *klp-19/spd-1 (RNAi)* 10.9µm ± 0.3µm, (n=38); *spd-1 (RNAi)* 11.7µm ± 0.2µm, (n=7); *hcp-6 (RNAi)* 11.4µm ± 0.2µm, (n=5), *hcp-6/ spd-1 (RNAi)* 11.5µm ± 0.2µm, (n=7), sem) (Suppl. Fig. 5A-D). At anaphase onset, spindles of *klp-19 (RNAi)* (14.4µm ± 0.4µm) and *klp-19/spd-1 (RNAi)* treated embryos (11.8µm ± 0.5µm) were significantly shorter in comparison to control (15.4µm ± 0.3µm), while spindles in *hcp-6 (RNAi)* (18.1µm ± 0.7µm) and *hcp-6/ spd-1 (RNAi)* (17.2µm ± 0.8µm) treated embryos where significantly longer than control (Suppl. Fig. 5A-D). Similar, 60s after Anaphase onset, spindles of *klp-19 (RNAi)* (19.2µm ± 0.5µm) and *klp-19/spd-1 (RNAi)* treated spindles (16.2µm ± 0.6µm) were significantly shorter in comparison to control (20.6µm ± 0.2µm), while spindles in *spd-1 (RNAi)* (24.5µm ± 0.9µm) treated embryos where significantly longer than control (Suppl. Fig. 5A-D). Spindles in *hcp-6 (RNAi)* (19.7µm ± 0.8µm) and *hcp-6/ spd-1 (RNAi)* (20.5µm ± 0.9µm) embryos were comparable to control. Quantification of chromosome segregation showed that chromosome segregation was significantly reduced in comparison to control (8.7µm ± 0.2µm) after depletion of KLP-19 (6.8µm ± 0.3µm), KLP-19/ SPD-1 (8.3µm ± 0.1µm), HCP-6 (5.1µm ± 0.2µm) and HCP-6/SPD-1 (5.7µm ± 0.4µm), while it was increased in embryos depleted of SPD-1 (14.1µm ± 0.8µm) (Suppl. Fig. 5A-D). Plotting the pole-to-pole distance over time revealed that HCP-6 depletion leads to a premature increase in spindle length prior to metaphase, with only a marginally increase in spindle length between meta- and anaphase (Suppl. Fig. 5C). In contrast, spindles of *klp-19 (RNAi)* embryos show decreased spindle length at the time of metaphase, with most increase in spindle length taking place during anaphase (Suppl. Fig. 5C).

When comparing spindle dynamics over time in embryos depleted of SPD-1, HCP-6/SPD-1 and KLP-19/SPD-1 (Suppl. Fig. 5D) we found clear differences in the dynamics, with KLP-19/SPD-1 depleted embryos showing shorter spindles.

As we observed a premature increase in spindle length in *hcp-6 (RNAi)* embryos prior to from Anaphase onset, we quantified the duration from NEBD to Anaphase onset to detect potential delays in cell cycle progression that could explain the premature spindle elongation. This showed that embryos depleted of HCP-6 had a delayed anaphase onset (223s ± 15, (n=8)) in comparison to control (166s ± 7, (n=7)) and *klp-19 (RNAi)* (185s ± 13, (n=9)), possibly caused by delays during congression or due to spindle checkpoint activity (Suppl. Fig. 5E). This suggests that the premature spindle elongation could be caused by pulling forces and a lack of stable kinetochore microtubule connections. Lastly, we wanted to test if depletion of HCP-6 had any effects on the microtubule overlap in the spindle. For this we quantified the distribution of SPD-1 GFP in the spindle midzone. We could not detect any changes in the microtubule overlap in response to HCP-6 depletion (Suppl. Fig. 5F). In summary this data suggests that while lagging chromosomes might affect spindle dynamics, the observed phenotypes after KLP-19 depletion differ significantly and are most likely not the result of chromosome misalignment alone but an indicator of its important role in the spindle midzone.

### KLP-19 regulates the microtubule overlap independently of the microtubule bundling protein SPD-1

PRC-1 has been suggested to recruit KIF4a to the spindle midzone in mammalian cells and in vitro, where both proteins interact to regulate the microtubule overlap (*36, 39, 40*). To assess the effect of *spd-1 (RNAi)* and *klp-19 (RNAi)* on the microtubule overlap in the midzone we used SHG and TP microscopy. Previous studies showed that SPD-1 localizes to and is required for the integrity of antiparallel MTs in the midzone (*57*). To further explore this, we studied the localization of SPD-1 in the one-cell *C. elegans* embryos. By correlating the localization of SPD-1-GFP with microtubule polarity, using a combination of 2-Photon and SHG microscopy, we found that SPD-1 indeed localizes to the region of antiparallel microtubules in *C. elegans* indicated by the overlapping profiles (Suppl. Fig 6A, note: the y-axis of the microtubule polarity has been inverted for easier viewing). This localization could suggest a potential role for SPD-1 in regulating overlap. To test this, we analyzed microtubule polarity in absence of SPD-1. As depletion of SPD-1 prevents the formation of a midzone in wild type embryos due to the extensive cortical pulling forces, we measured the microtubule polarity in spindles of *C. elegans* embryos co-depleted of SPD-1 and GPR-1/2. We could not detect any changes in the microtubule overlap in the central spindle, indicated by the overlapping polarity profiles, suggesting that although SPD-1 plays a key role in mechanically stabilizing the midzone, it is not required in determining the extend microtubule overlap when the pulling force is greatly reduced (Fig. 3A).

**Figure 3.**
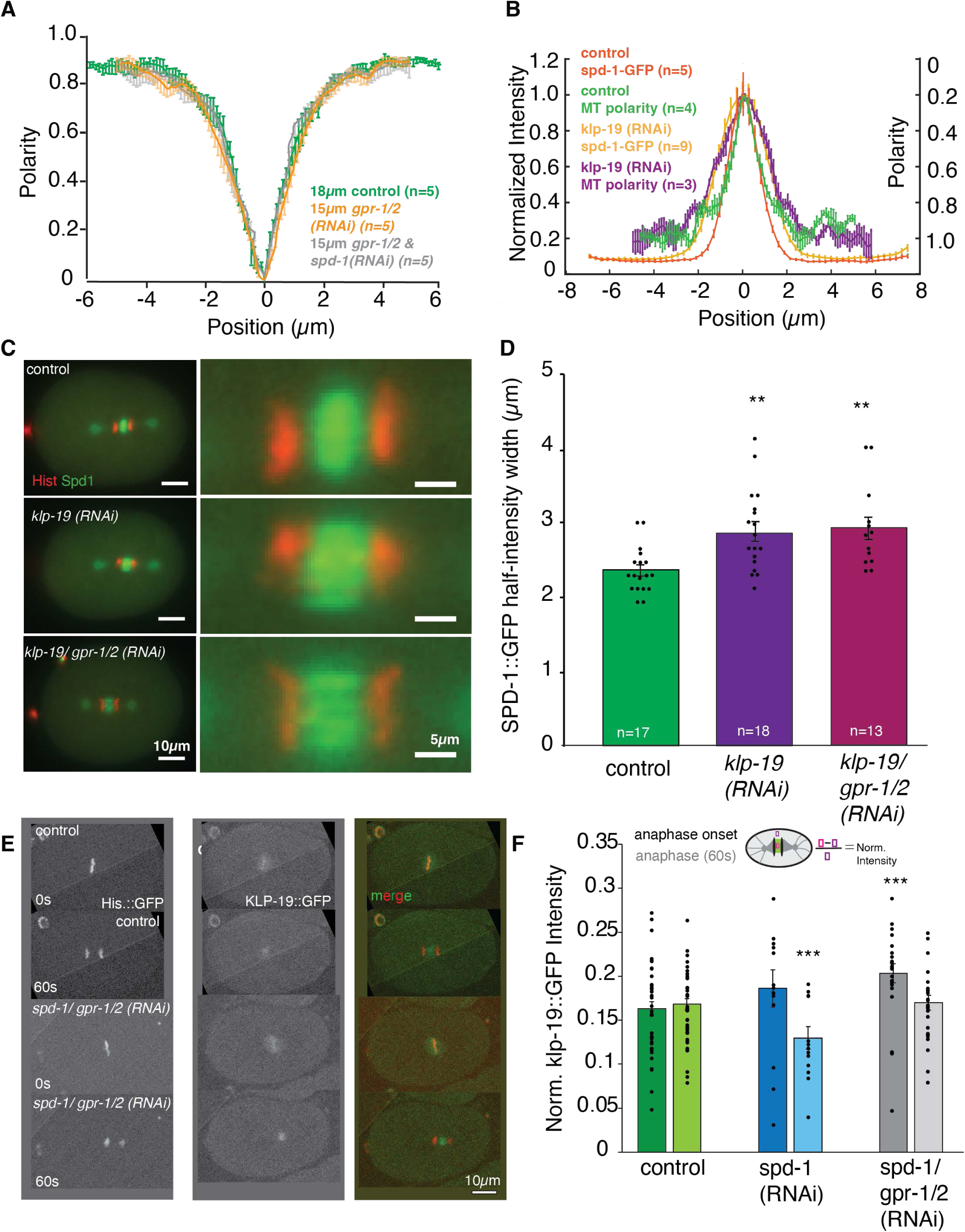
KLP-19 regulates the microtubule overlap length in the spindle midzone. **A.** Plot showing the polarity of microtubules along the spindle axis in control embryos (green), embryos depleted of pulling forces by *gpr-1/2 (RNAi)* (orange) and embryos after *spd-1/ gpr-1/2 (RNAi)* (grey). “0” is the spindle center **B.** Plot showing the normalized intensity (left y-axis) of SPD-1 GFP in control embryos (dark orange) along the spindle axis (0 is the spindle center on x axis) in anaphase and the microtubule polarity (right axis, green) along the spindle axis (right y-axis) in control embryos as well as SPD-1 GFP (bright orange) along the spindle axis in anaphase and the polarity of microtubules (purple) along the spindle axis in embryos after *klp-19 (RNAi)*. **C.** Left: Stills of anaphase spindles labeled with γ-tub::mCherry, histone::mCherry and SPD-1::GFP in control embryos (top) and embryos after *klp-19 (RNAi)* (bottom), Scale bar 10µm. Right: Zoom into the spindle midzone. Scale bar 5µm. **D.** Bar plot of the SPD-1 GFP half-intensity width in control (green), and *klp-19 (RNAi)* (purple). Error bars are sem. **E.** Stills of control embryos (left) and embryos after *spd-1/ gpr-1/2 (RNAi)* (right) in metaphase (top) and anaphase (bottom) expressing endogenously tagged KLP-19 GFP. **F.** Bar plot of KLP-19 GFP intensity at anaphase onset and 60s after anaphase onset in control embryos, embryos treated with *spd-1 (RNAi)* (blue) and *spd-1/ gpr-1/2 (RNAi)* (grey). Error Bars are sem. The significance of differences between control and RNAi conditions was determined by two-tailed Student’s *t*-tests (\*\*\*\**P* < 0.0001). Scale Bar 10µm.

Since the ratio of PRC1/KIF4A controls the extent of antiparallel microtubule overlap *in vitro* (*22, 41, 42*), we measured the localization of SPD-1 as well as the microtubule polarity in embryos depleted of KLP-19 (Fig. 3B). This showed a significant increase in SPD-1-GFP distribution in KLP-19-depleted embryos (Fig. 3C, D, Suppl. Fig. 6B), consistent with previous results (*22*). In agreement with this we found that the antiparallel microtubule region increases after KLP-19 depletion, indicated by an increase in width of the polarity profile measured by SHG and TP signals (Fig. 3B, Suppl. Fig. 6C, D). Next, we co-depleted KLP-19 and SPD-1 and quantified the profile of microtubule polarity. We found that the width of the antiparallel microtubule overlap region is increased in these embryos in comparison to wildtype and comparable to the KLP-19 depletion (Suppl. Fig 6C, D). This data shows that KLP-19 regulates the size of the antiparallel microtubule overlap and thus narrows the distribution of SPD-1, confirming a role for the *C. elegans* KLP-19 in the spindle midzone in addition to its previously reported role during congression (*58*).

Based on previous publications we assumed that depletion of SPD-1 would affect the localization of KLP-19 to the midzone in *C. elegans*. However, this is inconsistent with the observed effect on microtubule overlap and the different phenotypes upon double depletion.

To follow up on this observation we generated a worm strain carrying endogenously GFP tagged KLP-19 using CRISPR technology. This strain allowed us to monitor the localization of KLP-19 throughout mitosis. We found that KLP-19 is initially enriched in the pro-nuclei and localizes to chromosomes (Fig. 3E). Upon Nuclear envelope breakdown (NEBD) KLP-19 localizes to chromosomes as well as spindle microtubules but could not be detected on centrosomes or astral microtubules. At anaphase onset KLP-19 transitions from the chromosomes to the spindle midzone (Fig 3E). To determine the dependency of KLP-19 midzone localization on SPD-1 we monitored the localization of KLP-19 GFP during mitosis in *spd-1 (RNAi)* and *spd-1/ gpr-1/ 2 (RNAi)* embryos. The KLP-19 GFP signal in the spindle midzone was mainly lost after *spd-1 (RNAi),* due to premature spindle rupture that prevented the formation of a midzone (Fig. 3F).

However, the localization of KLP-19 to the spindle midzone in absence of pulling forces, preventing premature spindle breakage by *spd-1/ gpr-1/ 2 (RNAi,)* was unchanged, suggesting that KLP-19 localizes to the spindle midzone independently of SPD-1 in *C. elegans* (Fig. 3F). Likewise, localization to chromosomes was unchanged in either RNAi treatment (Fig 3F).

To further establish how the localization of KLP-19 to the spindle midzone is regulated we depleted the kinetochore proteins BUB-1, HCP-3, HCP-4, NDC-80 and AIR-2. We found that depletion of AIR-2, NDC-80 and HCP-3 increased the amount of KLP-19 on chromosomes in metaphase (Suppl Fig. 7), while depletion of BUB-1 led to a reduction of KLP-19. In anaphase, depletion of AIR-2 and BUB-1 significantly reduced KLP-19 in the midzone. We also detected an increase of KLP-19 in the midzone after depletion of NDC-80, however as chromosome segregation fails in these embryos it is very likely that the increase reflects the amount of KLP-19 on the chromosomes and not in the midzone. These data suggest that the localization of KLP-19 to the spindle midzone is dependent on AIR-2 and BUB-1, as previously reported (69,70).

### KLP-19 is a processive Microtubule motor in C. elegans and does not interact with SPD-1

The effects of KLP-19 and SPD-1 on the overlap of midzone microtubules prompted us to ask whether KLP-19 is functioning as a processive microtubule motor in mitotic *C. elegans* zygotes. To address this, we developed an *ex vivo* assay for motor activity that is based on a single-cell, single-molecule pull-down (sc-SiMPull) assay (Fig. 4A) (*62*). Briefly, we fabricated a microfluidic device consisting of a coverslip and polydimethylsiloxane (PDMS). In a typical sc-SiMPull assay, the coverslip surface would be coated with antibodies for single-molecule pull-down, but here, we attached pre-polymerized, paclitaxel-stabilized, biotinylated, and fluorescently labeled microtubules instead. A zygote carrying endogenously tagged KLP-19 was then introduced into the channel and lysed during anaphase using an infrared pulsed laser, which creates a cavitation bubble and shockwave to disrupt the cell membrane. Starting within 3-5 seconds after lysis, we observed the samples using total internal reflection (TIRF) microscopy. Strikingly, we found that cell-derived KLP-19::GFP molecules bound and processively moved on the immobilized microtubules (Fig. 4Bii-iv, Movie 6, 7). Using single particle tracking, we measured an average speed of approximately 0.2 μm/sec (Fig. 4C, Suppl Fig. 8A) with a run length of about 2 μm (Fig. 4D, Suppl Fig. 8B), similar to the *Xenopus* Kif4a homolog, Xklp-1 that was measured as a purified protein (*40*). Because human KIF4a exists as a homodimer (*38, 40, 63*), we were curious to determine the stoichiometry of KLP-19. Using conventional sc-SiMPull, we captured endogenous KLP-19::GFP in a microfluidic device coated with anti-GFP antibodies (Suppl Fig. 8C-left). We then used single-molecule photobleaching to count the number of KLP-19 molecules in each complex (Suppl Fig. 8C-right). KLP-19 formed homodimers, similar to human Kif4a (Suppl. Figure 8D). Returning to our data from the motility experiment, we then asked whether oligomerization of KLP-19 might occur on microtubules and could influence its motility. However, the brightness of moving KLP-19 particles fell within a relatively narrow range and was not correlated with the speed of the particles (Suppl Fig. 8E), suggesting that higher-order oligomerization of KLP-19 homodimers did not occur (and did not influence KLP-19 speed) in this *ex vivo* assay. Altogether these data showed that KLP-9 is a dimeric motor protein that can processively move on microtubules.

**Figure 4.**
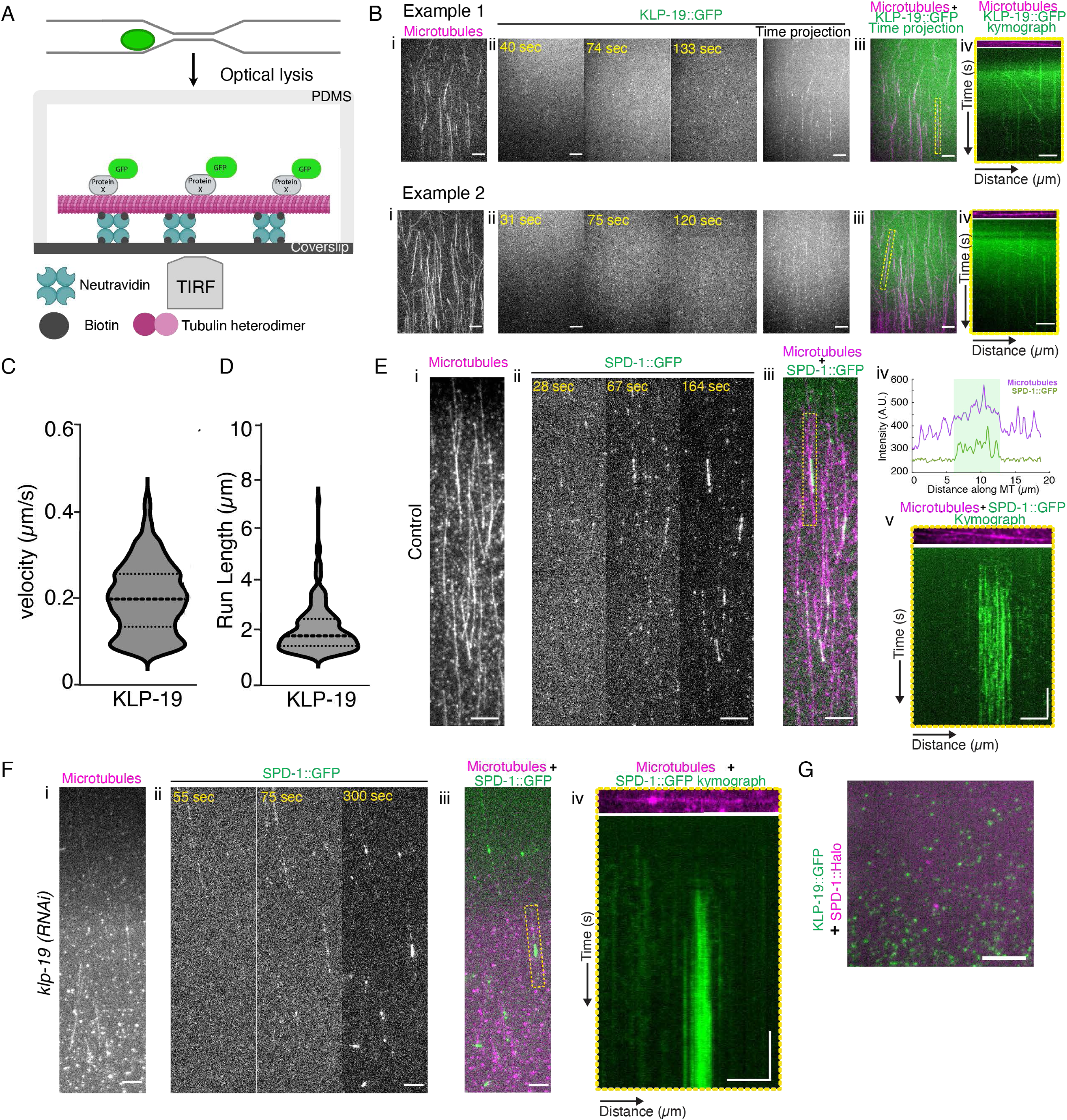
KLP-19 is a processive motor and does not interact with SPD-1. **A**. Schematic of the *ex vivo* motility assay. Briefly, an embryo expressing an endogenously tagged protein of interest is trapped in a PDMS microfluidic device. The embryo is then lysed using an infrared pulsed laser. The fluorescently tagged protein diffuses from cell lysate and lands on the surface of the coverslip, which is coated with *in vitro* polymerized microtubules, tethered to the coverslip through biotin-neutravidin binding. The binding and movement of the protein on microtubules are captured by a TIRF microscope. **B**. (i) Still TIRF image from a time-lapse movie showing immobilized fluorescently labeled microtubules. (ii) TIRF montage images from a timelapse movie showing KLP-19::GFP single molecules that appear on the surface of the coverslip. Maximum-intensity time projections showing that over time KLP-19::GFP landing events are strongly enriched along microtubules. (iii) Overlay of the KLP::GFP time projection and microtubule images. (iv) Top: still TIRF image of the microtubule highlighted in yellow in (iii). Bottom: a kymograph showing single KLP-19::GFP particles moving along the microtubule. All horizontal scale bars represent 5μm, vertical scale bar represents 50 seconds. **C**. Plot showing the velocity of KLP-19::GFP single particles. For clarity, only particles with a velocity >0.075 µm/s are shown here; see Figure S8A for all data. **D**. Plot showing the run lengths of KLP-19::GFP single particles. For clarity, only particles with a run length >1.25 µm are shown here; see Suppl. Fig. 8B for all data. **E**. (i) Still TIRF image showing immobilized labeled microtubules. (ii) TIRF images from a timelapse movie of SPD-1::GFP binding microtubules and concentrating at a region of dense microtubules in control embryos. (iii) Overlay of the SPD-1::GFP image at 164 seconds after lysis and the microtubule image. (iv) Line scans showing tubulin (magenta) and SPD-1::GFP (green) intensity along the length of microtubule shown in Figure 4E. (v) Top: still image of a single microtubule from the area in (iii) highlighted in yellow. Bottom: Kymograph of SPD-1::GFP particles. **F**. (i) Still TIRF image showing immobilized labeled microtubules. (ii) TIRF images from a timelapse movie of SPD-1::GFP binding microtubules and concentrating at a region of dense microtubules in *klp-19 RNAi* embryos. (iii) Overlay of the SPD-1::GFP image at 300 seconds after lysis and the microtubule image. (iv) Top: still image of a single microtubule from the area in (iii) highlighted in yellow. Bottom: Kymograph of SPD-1::GFP particles. **G**. Average intensity projection of 50 frames from a sc-SiMPull experiment showing captured KLP-19::GFP molecules (green) that are not colocalized with SPD-1::HaloTag (magenta). All horizontal scale bars represent 5μm, vertical scale bars represent 50 seconds.

We further examined the behavior of SPD-1 *ex vivo* on immobilized microtubules. We observed that SPD-1 bound to microtubules and concentrated at regions of denser and brighter tubulin signal, which we suspect to be regions where two immobilized microtubules overlapped (Fig. 4Ei-iv, Movie 8). We analyzed single SPD-1 particles and found that unlike KLP-19 (Fig. 4B, C), SPD-1 did not move along microtubules (Fig. 4Ev). Furthermore, when we repeated these experiments after depleting KLP-19 by RNAi, SPD-1 still concentrated in areas where microtubules putatively overlapped (Fig. 4F). Together, these data suggest that SPD-1 is not transported by KLP-19 as had been suggested for mammalian PCR1 and KIF4a (*26*). These observations prompted us to ask whether SPD-1 and KLP-19 interact. To test this, we constructed a strain carrying endogenously tagged KLP-19::GFP and SPD-1::HaloTag, and used sc-SiMPull (*62*) to look for an interaction between these two proteins in anaphase zygotes. We counted a total of 70,670 KLP-19::GFP molecules from 7 anaphase zygotes, but found only 39 molecules (0.05%) that appeared to be associated with SPD-1::HaloTag (Fig. 4G) – well below the background levels we have previously reported using this assay (*62, 64*). Thus, we could find no evidence for a complex between KLP-19 and SPD-1 in anaphase *C. elegans* zygotes, even though the mammalian homologs of these proteins have been reported to interact *in vitro* and in cells (*22, 26, 36, 37*). Although this is a negative result and we cannot formally exclude the possibility that KLP-19 and SPD-1 interact transiently, we favor the hypothesis that interactions between these proteins *in vivo* are indirect and mediated by midzone microtubules.

### KLP-19 regulates microtubule dynamics in the spindle midzone

Since KLP-19 regulates microtubule overlap in the spindle midzone and had previously been reported to negatively regulate microtubule dynamics we wanted to quantify its effect on microtubules and the midzone in *C. elegans* in more detail. Analysis of microtubule intensity in the spindle midzone after *klp-19 (RNAi)* (0.81A.U. ± 0.07A.U., (n=19), sem) showed a significant increase in intensity in comparison to control (0.48A.U. ± 0.04A.U, (n=29)), suggesting a potential increase in microtubule number or density (Fig 5A). Also, co-depletion of KLP-19 and GPR-1/2 (0.57A.U. ± 0.05A.U., (n=14), used as the control for a following FRAP experiment Fig, 5B) as well as KLP-19 and SPD-1 (0.75A.U. ± 0.07A.U, (n=16)) lead to increased microtubule intensities in the midzone, while *spd-1 (RNAi)* leads to a decrease in microtubule intensity (0.30A.U. ± 0.06A.U, (n=18)). Depletion of GPR-1/2 did not affect the intensity (0.41A.U. ± 0.04A.U, (n=11)). Interestingly, we found that co-depletion of SPD-1 and GPR-1/2 leads to an increase in microtubule intensity (0.73A.U. ± 0.06A.U, (n=8)). Based on the proposed function of KIF4A in stalling microtubule plus-end growth, we speculated that depletion of KLP-19 could affect microtubule dynamics in *C. elegans.* We used Fluorescence Recovery After Photobleaching (FRAP) to determine the turn-over of microtubules in absence of KLP-19. As the mitotic spindle moves and oscillates significantly due to the cortical pulling forces in anaphase, which makes FRAP experiments difficult, we quantified and compared the effect on microtubule turn-over in the center of the spindle midzone during anaphase after depletion of either GPR-1/2 alone or KLP-19 together with GPR-1/2 (to reduce spindle motion and oscillations). The data did not show a significant difference between the fraction of microtubule recovery after *klp-19 (RNAi).* However, we did observe a trend towards a decreased microtubule recovery after KLP-19 depletion (0.23 ± 0.04, (n=23) vs 0.34 ± 0.06, (n=10) in control), suggesting a possible reduction in microtubule turn-over (Fig. 5B).

**Figure 5.**
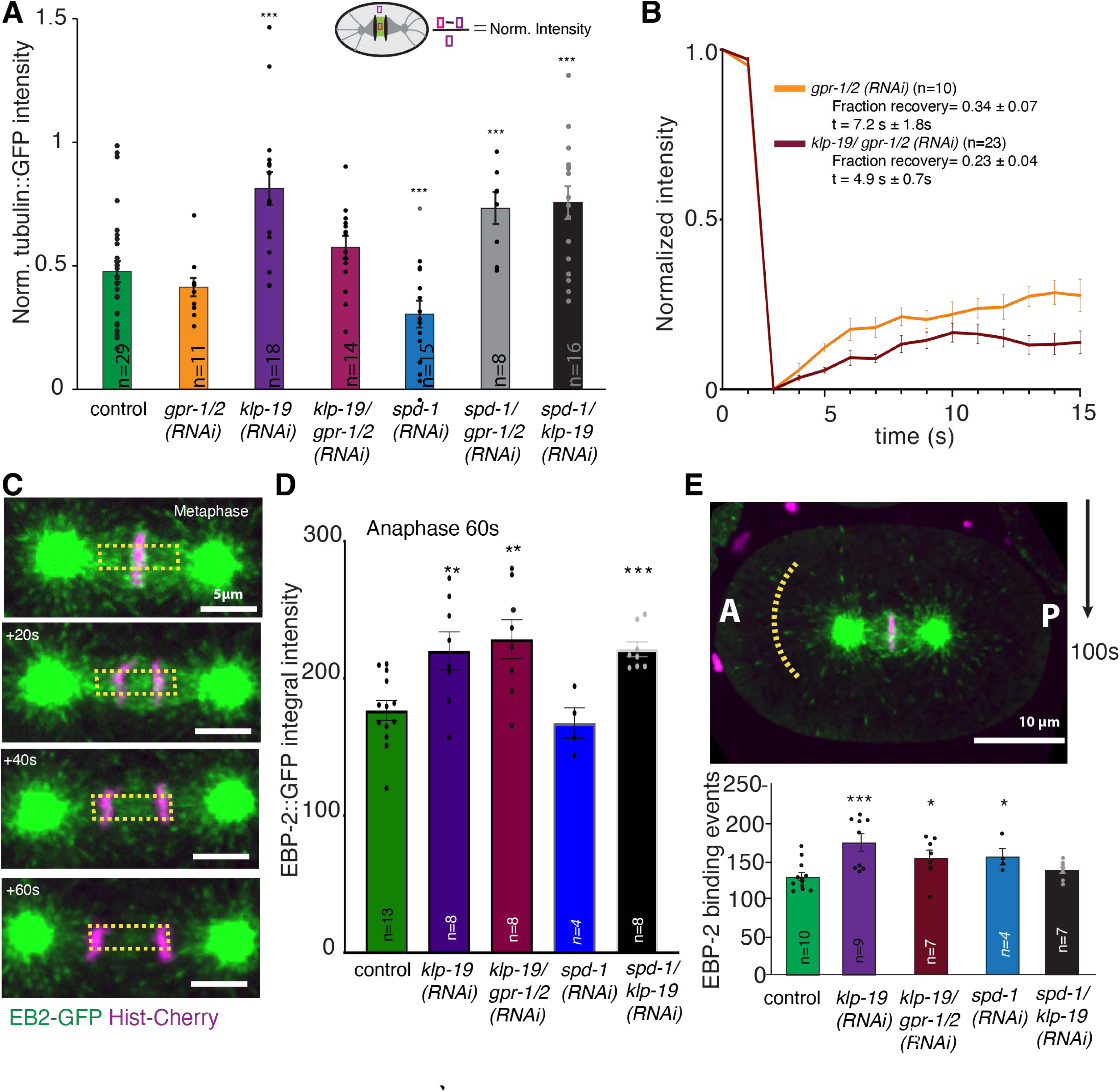
KLP-19 affects microtubule dynamics. **A.** Plot of the normalized α-tubulin::GFP intensity in the spindle midzone 60s after anaphase onset in control embryos and after *gpr-1/2 (RNAi)*, *klp-19 (RNAi)*, *klp-19/gpr-1/2, spd-1 (RNAi)*, *spd-1/gpr-1/2 (RNAi)*, and *spd-1/klp-19 (RNAi).* **B**. Plot of Microtubule fluorescence recovery over time after photobleaching (FRAP) in the spindle midzone in anaphase in control embryos and embryos after *gpr-1/2 (RNAi)* and *klp-19/gpr-1/2 (RNAi)*. **C.** Stills of a *C. elegans* embryo in anaphase expressing EBP-2::GFP and Histone::mCherry. Yellow box indicates the area used for intensity measurements shown in the corresponding plots in D. Scale Bar 5µm. **D.** Bar plot showing the normalized intensity of EBP-2::GFP signal in the spindle midzone in control embryos and after *klp-19 (RNAi), klp-19/ gpr-1/2 (RNAi), spd-1 (RNAi)* and *spd-1/ klp-19 (RNAi)* 60s after anaphase onset. **E.** Top: Still of an embryo expressing EBP-2::GFP and mCherry Histone. Anterior and posterior pole are indicated. Yellow half-circle on the posterior indicates the region of EBP-2 signal quantification. Scale Bar 10µm. Bottom: Bar plot showing the quantification of EBP-2 comets at the yellow line over 100s. 4.5µm length segment of the line was analyses for binding events. Color code: control embryos = green, *gpr-1/2 (RNAi)* = orange, *klp-19 (RNAi)* = purple, *klp-19/ gpr-1/2 (RNAi) =* maroon, *spd-1 (RNAi)* = blue, *spd-1/ gpr-1/2 (RNAi)* = grey, *klp-19/ spd-1 (RNAi)* = black. Error Bars are sem. The significance of differences between control and RNAi conditions was determined by two-tailed Student’s *t*-tests (\*\*\*\**P* < 0.0001).

To determine effects on microtubule growth we imaged embryos expressing GFP-tagged EBP-2, which localizes to growing microtubule plus-ends. Quantification of the EBP-2 signal showed an increase in EBP-2 integral intensity in the midzone upon *klp-19 (RNAi)* (220.1 A.U. ± 13.9, n=8, sem) in comparison to control (176.6A.U. ± 7.2, n=13), suggesting increased microtubule polymerization (Fig. 5C, D). In agreement with that, co-depletion of KLP-19 and GPR-1/2 (228.4A.U. ± 14.23A.U, n=8) as well as KLP-19 and SPD-1 (221.1A.U. ± 5.4A.U, n=8) led to increased EBP-2 integral intensity in the midzone, while *spd-1 (RNAi)* (167.5A.U. ± 10.98A.U, n=4) did not affect the intensity of the EBP-2 signal. This result is consistent with the proposed stalling of microtubule growth by KLP-19 (*65*). The effect on microtubule dynamics could be restricted to the midzone or affect global microtubule dynamics. Quantification of EBP-2 binding events on astral microtubules in anaphase showed an increase in embryos depleted of KLP-19 (178 ± 11.42 events) in comparison to control (132.2±5.6 events) (Fig. 5E). As our data showed an increase in tubulin intensity and an increase in EBP-2 binding events after *klp-19 (RNAi)* we conclude that KLP-19 globally regulates microtubule dynamics during mitosis in *C. elegans,* possibly by reducing polymerization and eventually promoting turn-over by inducing depolymerization.

### Depletion of KLP-19 affects the architecture of the spindle midzone

To quantify the effect of KLP-19 depletion on microtubule length in the spindle midzone we used 3D tomography to analyze the microtubule arrangement in the spindle midzone in response to *klp-19 (RNAi)*. (Fig 6A, B, Suppl. Fig. 9). Quantification of microtubule number, length and the interaction between microtubules in the midzone in the tomographic reconstructions revealed an increase in the average microtubule number from 3.1 microtubule/µm^3^ in control to 4.3 microtubule/µm^3^ and an increase in average microtubule length from 0.88 µm ± 0.14 µm (STD) in control to 1.15 µm ± 0.03 µm (STD) after *klp-19 (RNAi)*. In addition, we quantified the average distance between microtubules in the reconstructions, which was 71nm ± 2nm (STD) in control and 62nm ± 2nm (STD) after KLP-19 depletion. The microtubule overlap length changed from 0.42 µm ± 0.02 µm (STD) in control to 0.50 µm ± 0.07 µm (STD) after *klp-19 (RNAi)* (Fig. 6C, D, Suppl. Fig.9).

**Figure 6.**
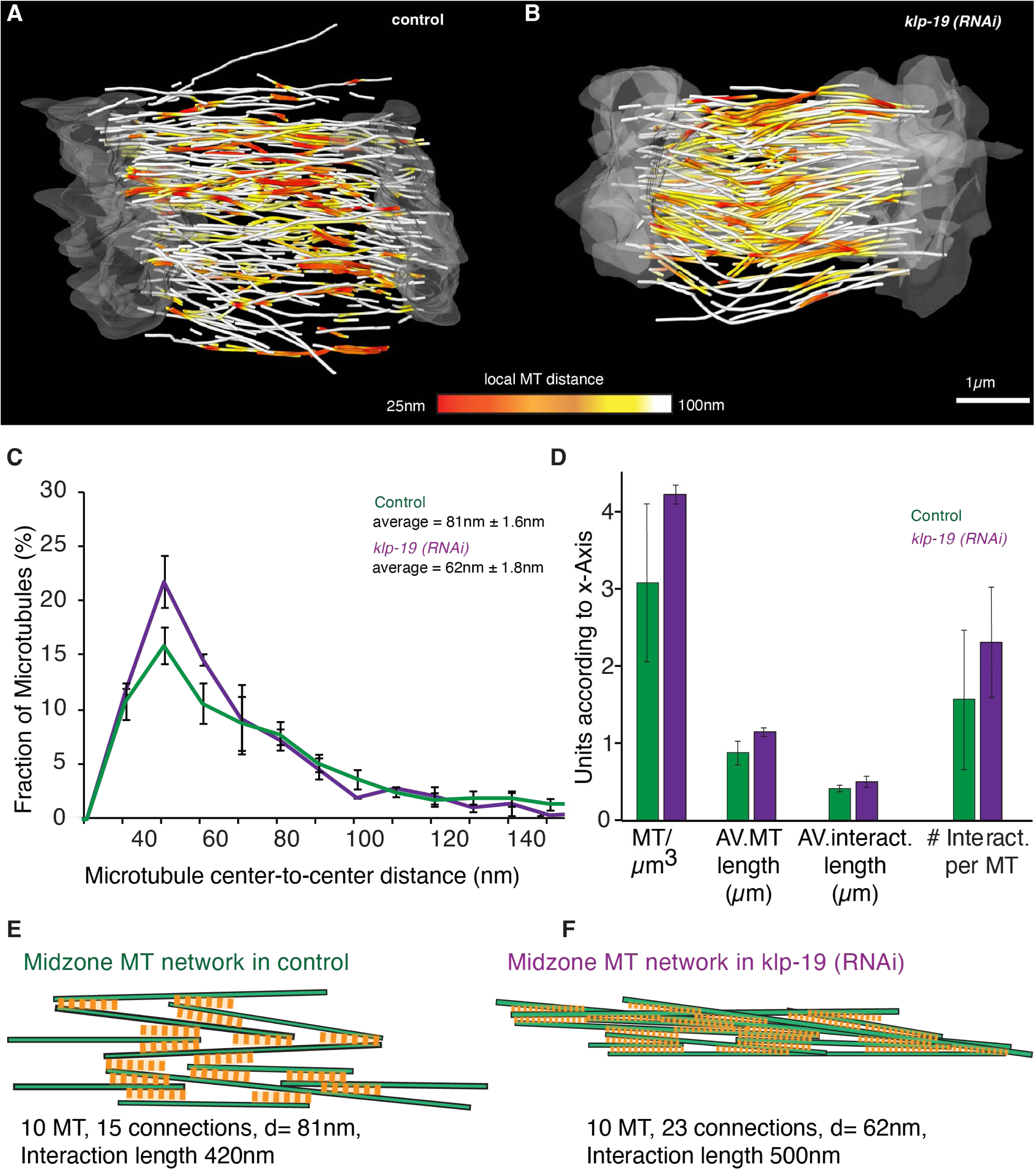
KLP-19 depletion leads to increased microtubule length and interactions. **A.** Representative 3D tomographic reconstruction obtained by electron tomography of control (left) and *klp-19 (RNAi)* (right) embryos showing only the microtubules in the spindle midzone (see also Suppl. Fig. 9). Microtubules are color coded according to the nearest local distance (center to center) to a neighboring microtubule, with red being 25nm and white larger than 100nm. Scale bar is 1µm **B.** Plot of the fraction of microtubule interactions within a defined microtubule center-to-center distance. Average microtubule distance is provided in the plot. Error Bars are STD **C.** Bar plot of different average parameters for microtubules in control (green) and *klp-19 (RNAi)-*treated embryos (purple). Error Bars are STD **E.** Cartoon visualizing the interactions of microtubules in the mitotic spindle midzone in control embryos during anaphase, based on data retrieved from the 3D reconstructions. Parameters used for the cartoon are indicated. **F.** Cartoon visualizing the interactions of microtubules in the mitotic spindle midzone in embryos after *klp-19 (RNAi)* during anaphase, based on data retrieved from the 3D reconstructions.

These data suggests that KLP-19 is involved in regulating the microtubule length and interaction and thus the overlap in the central spindle. Depletion of KLP-19 increases the microtubule overlap and interaction, leading to a more connected microtubule network in the spindle midzone (Fig. 6E, F). A more connected microtubule network could make it harder to break the spindles as more microtubule interactions would need to be broken. In addition, increasing the length of microtubules could provide more space for cross-linkers or motor proteins to bind and thus increase the amount of interaction.

## Discussion

In this study we addressed the role of the KIF4a homolog KLP-19 in the spindle midzone during anaphase in the *C. elegans* one-cell embryo. Numerous conserved proteins are associated with the midzone, coordinating its assembly and function. The microtubule cross-linker PRC1 plays a critical role in midzone stability in mammalian cells. In agreement with this, depletion of the *C. elegans* homolog, SPD-1, leads to spindle rupture during mitosis in response to pulling forces.

Previous data has proposed that both PRC1 and Kif4a are involved in the regulation of microtubule overlap in the spindle midzone (*29, 41, 66, 67*). Our study demonstrates that the overlap of microtubules in the midzone remains constant during anaphase and is independent of pulling forces. We find that the size of the microtubule overlap in *C.elegans* is dependent on KLP-19, as the overlap increases upon *klp-19 (RNAi),* but independent of SPD-1. Our data suggests that KLP-19 affects microtubule dynamics, most likely by stalling microtubule growth and possibly by promoting turn-over, which is in agreement with previously reported roles of the mammalian homolog KIF4a (*36–38*). Loss of KLP-19 thus leads to increased microtubule growth and length in the midzone and with that an increase in microtubule interactions.

Аlthough some previous publications suggested that PRC1 recruits Kif4a (*22, 26, 36–38, 40*), we could not detect a decrease in KLP-19 signal in the midzone after depletion of SPD-1 in *C. elegans* embryos. In agreement with that our single cell pull-down experiments did not detect an interaction between KLP-19 and SPD-1, suggesting that these proteins do not directly interact and that localization of KLP-19 to the midzone is independent of SPD-1. This is consistent with other previous studies showing that compacted midzone organization is driven by Kif4a and not dependent on its known interaction with PRC1 via the C-terminal tail domain (*36*).

Previous publications found that depletion of PRC1 prevented the localization of KIF4a to the midzone in mammalian cells (*22, 28, 29, 44*). Based on our results we speculate, that in both *C. elegans* and mammalian cells, cortical pulling forces (*68*) might prevent the formation of a midzone upon PRC1 depletion. This could explain the reported hypersegregation phenotype after PRC1 depletion (*44*). An absent or significantly weakened microtubule midzone after PRC1 depletion could thus prevent the localization of KIF4a to the midzone. In situations where pulling forces are absent, i.e. in *C. elegans* after depletion of GPR-1/2 a microtubule based midzone still forms, even after depletion of SPD1, allowing KLP-19 and other midzone proteins to localize to the midzone. This suggests that KLP-19 might not rely on SPD-1 for midzone localization but rather on the presence of a microtubule network in the midzone and that this could be similar in mammalian cells.

Further analysis showed that the localization of KLP-19 to the midzone is dependent on BUB-1 as well as AIR-2. This is in agreement with previously published work showing that KLP-19 requires BUB-1 to localize between the bivalents in *C. elegans* meiosis (*69*). As the localization of BUB-1 is dependent on AIR-2 (*69, 70*) it is likely that AIR-2 could prevent the localization of KLP-19 to the midzone by inhibiting the localization of BUB-1. However, in mammalian cells Aurora B phosphorylation of KIF4A has been shown to activate its microtubule-stimulated ATPase activity and enhances KIF4A binding to PRC1 and the antiparallel microtubule overlap (*37*,*71*) suggesting that there could also be a direct interaction of KLP-19 with AIR-2. Further investigations would be required to determine the mechanism of KLP-19 translocation to the spindle midzone.

KLP-19 has been shown to play an important role during chromosome congression and the generation of polar ejection forces (*58*). Thus, KLP-19 has been described as a chromokinesin, sliding chromosomes towards the plus-end of microtubules. Our study confirmed the role of KLP-19 during congression. However, while we can not exclude that KLP-19 actively transports chromosomes, based on our data and KLP-19’s effect on microtubule dynamics, it is also possible that chromosome-bound KLP-19 directly affects microtubule dynamics at the chromosomes. Regulating microtubule dynamics could then affect the forces that microtubules generate on chromosomes during congression, thus contributing to chromosome alignment. Therefore, it would be interesting to determine the detailed role of KLP-19 during congression in the future.

The spindle midzone serves as a mechanical link between the two spindle halves, experiencing tension from cortical force generators crucial for spindle positioning while actively contributing to chromosome segregation by promoting spindle elongation. This mechanical balance between stability and adaptability is essential for ensuring the proper segregation of chromosomes, not only in *C. elegans* embryos but also in mammalian cells, which experience pulling forces as well (*68, 72, 73*). The midzone is a microtubule network held together by motor proteins and cross-linkers, forming dynamic, reversible bonds that allow for rearrangements and plasticity throughout anaphase. Stability within this network depends on the equilibrium between connection breakage and reformation. The observed stereotypical size of the midzone, with or without cortical pulling forces, suggests a tight regulation of the overlap zone throughout anaphase, indicating an inherent balance between microtubule dynamics and interaction of proteins with midzone microtubules.

It is plausible that the length of microtubules plays a pivotal role in midzone stability, with longer microtubules enhancing stability but potentially impeding chromosome segregation through increased interactions.

Our data suggest that the increased overlap and interaction of microtubules in the midzone, resulting from KLP-19 depletion, compromises the mechanics of chromosome segregation. The absence of KLP-19 results in a more interconnected midzone network, which could explain the observed reduced rates of chromosome and pole separation during mitosis.

We can not unambiguously rule out that failure to properly align chromosomes and the resulting lagging chromosomal material could also lead to some of the observed effects on spindle dynamics, such as slow chromosome segregation and pole separation rates as well as preventing spindle rupture in absence of SPD-1. However, several observations argue in favor of KLP-19 actively changing the midzone cytoskeleton network and thus affecting spindle dynamics.

Our analysis of HCP-6 mutants generating excessive lagging chromosomes reveals drastic changes in spindle dynamics but fails to reproduce the reduced spindle size and chromosome segregation rates during mitosis observed in *klp-19 (RNAi)*. Similarly, depletion of the kinetochore protein KNL-1 was shown to lead to errors in chromosome congression as well as to abolish chromosome segregation (*6*). While chromosomes do not segregate, a rapid and premature pole separation can be observed in absence of KNL-1, comparable to the spindle dynamics upon *hcp-6 (RNAi)*.

This data shows that defects in chromosome alignment or failure to establish correct kinetochore microtubule connections does affect spindle dynamics, but usually leads to increased rates of pole separation rates and distance. This suggests that the observed spindle dynamics in absence of KLP-19 are not only caused by lagging chromosomes. Instead, KLP-19 RNAi results in a global rearrangement of the spindle and leads to a significant reduction of the spindle size, microtubule overlap, growth rate, and stability. Furthermore, the increase of microtubule interactions after klp-19 (RNAi) could also contribute to lagging of chromosomes and exacerbation of fragmented extrachromosomal material.

To summarize, in this study, we present evidence of the stability and precise regulation of microtubule overlap within the midzone, highlighting its significant role in chromosome segregation. For the first time, using 3D electron microscopy, we demonstrate *in vivo* that KLP-19, the *C. elegans* homolog of the human KIF4a, affects microtubule length within the spindle midzone. Our data underscores the importance of regulating microtubule dynamics and length to ensure proper mechanics during chromosome segregation, which is essential for accurate genetic distribution during cell division.

Additionally, we have developed a novel in vitro assay that enables us to observe endogenous microtubule-associated proteins from single-cell, staged *C. elegans* embryos. This has allowed us to visualize endogenous microtubule binding proteins, such a KLP-19 and SPD-1 at single-molecule resolution, paving the way for future research into motor proteins and potentially other protein behaviors.

## Materials and Methods

### CRISPR/Cas9 injections

Klp19-GFP tagged line was created via CRISPR/Cas9 mediated genome engineering using the self-excising cassette (SEC) method to facilitate screening and the guide RNA targeting sequence 5’ CAAGCGAAAGAGTCGTCGAA 3’ (Dickinson et al., 2013, 2015). Targeting sequence was cloned into pDD162 plasmid by using NEB’s Q5 Site-Directed Mutagenesis Kit to insert the targeting sequence into our Cas9-sgRNA construct (<u>Addgene #47549</u>) by using forward primer 5’-(CAAGCGAAAGAGTCGTCGAA)GTTTTAGAGCTAGAAATAGCAAGT-3’, and reverse primer 5’-CAAGACATCTCGCAATAGG-3’. The repair templates were generated by first cloning homology arms, synthesized by PCR using genomic DNA as a template, into plasmid: pDD282 with C-terminal GFP Tag with FLAG and flexible linker. Worm injections and transgenics selection was done according to the above cited protocol. The strain SR52 was created, expressing endogenously labeled KLP-19::GFP.

Spd-1::HaloTag tagged line was created in the same way as described above. The targeting sequence was cloned into pDD315 by using forward primer 5’ATACTATACGAAGTTATT TTTCAGGGAGCCGGCTGAATAATTTCATCATTCCAATTT 3’ and reverse primer 5’ACAGCTATGACCATGTTACCGGTATCGATTTCTTTCCAGCAAAAACCGTGAC 3’.

Injections were done into the OD3 line and transgenics selection was done according to the above cited protocol. The strains APL524 (Roller) and APL525 (excised) were created, expressing endogenously labeled SPD-1::HaloTag. Excision was done by maintaining worms in 34°C for 4 hours.

### C. elegans strains

MAS91 (unc-119(ed3) III; ItIs37[pAA64; pie-1::mCherry::HIS58]; ruIs57[pie-1::GFP:: α-tubulin+unc-119(+)]) was used for experiments of fluorescence imaging, laser ablation, and fluorescence recovery after photobleaching. MAS91 was a gift from Martin Srayko. ANA72 (adeIs1 [mex-5::spd-1::GFP + unc-119(+)] II. ltIs37 [pie-1p::mCherry::his-58 + unc-119(+)] IV.) was a gift from Marie Delattre. ANA72 was used to quantify the localization of SPD-1.

*wow47 [ebp-2::gfp::3xflag)] II; zif-1(gk117) III; wyEx9745]* was a gift from Kang Shen. OD305 (*unc-119(ed3) III; ltIs138 [pJD14; pie-1/KNL-1::GFP ; unc-119 (+)], unc-119(ed3) III; ltIs37 [pAA64; pie-1/mCherry::his-58; unc-119 (+)] IV)* was a gift from Karen Oegema and Arshad Desai.

Wild-type (N2) C. elegans embryos were used for the preparation of electron tomography. All strains were maintained at 16°C and fed on OP50 bacteria on nematode growth medium plates (Brenner, 1974).

OD3 (unc-119(ed3) III; ltls24.ltls24[(pAZ132) pie-1p::GFP::tba-2+unc-119(+)) was obtained from CGC.

SR61 was created by mating APL524 (SPD-1 HaloTag) into SR52 (KLP-19 GFP) and subsequent heatshock of 4h at 34 °C.

### RNAi

RNAi experiments were performed by feeding [Fire, 1998, Potent and specific genetic interference by double-stranded RNA in Caenorhabditis elegans]. RNAi feeding was performed on NGM plates containing 50 mg/mL ampicillin and 1 mM IPTG (Isopropyl b-D-thiogalactopyranoside) and seeded with the HT115(DE3) bacterial strain containing the desired target sequence.

Feeding clones for GPR-1/2, SPD-1, KLP-7 and KLP-19 were obtained from the RNAi library (*74*). L4 larvae were transferred to plates seeded with bacteria producing dsRNA and grown for 48 hr at 25C prior to analysis.

The feeding clone for AIR-2 (B0207.4) was purchased from Horizon Discovery. Feeding clones for BUB-1 (CUUkp3300H113Q), HCP-3 (DFCIp3320C1010038D), HCP-4 (CUUkp3300L171Q) and NDC-80 (DFClp3320H0110028D) were purchased from Source Bioscience. The Feeding clone for ZYG-1 was a gift from Kevin O’Connell (*75*).

RNAi in Fig 4F was done by injection. 1.5kb was amplified from N2 cDNA using primers containing T7 promoter overhang at both ends. PCR products were purified, transcribed using Promega T7 RiboMax Express kit (Cat. No. P1700) and annealed to make dsRNA. dsRNA was purified and injected into young adults at 1ug/uL concentration. Worms were imaged 24hrs post injection.

### Spinning disk confocal fluorescence imaging

Embryos for live-imaging experiments were obtained by dissecting gravid adult hermaphrodites in M9 buffer (42 mM Na_2_HPO4, 22 mM KH_2_PO_4_, 86 mM NaCl, and 1 mM MgSO_4_). One-cell embryos were mounted on slides with 2% agarose pad, overlaid with a 22 × 22-mm coverslip, and imaged at room temperature on a widefield fluorescence microscope (ECLIPSE Ti2; Nikon) equipped with a CFI Apo TIRF 60× 1.45 NA oil immersion lens and a CCD camera (iXon 897 Ultra EMCCDs) or on a 3i VIVO spinning-disc confocal microscope (Axio Examiner.Z1; Zeiss) equipped with Zeiss Plan-Apochromat 63x/1.40 oil microscope objective, 6 laser lines and a Hamamatsu ORCA-Flash4.0 scientific CMOS camera (Hamamatsu) for detection. The microscope was controlled by Nikon NIS-Elements software (Nikon). Imaging was initiated in one-cell embryos before pronuclear meeting and was terminated 3 min after anaphase onset.

EBP-2::GFP movies acquired with a spinning-disc confocal microscope. Image processing was done with ImageJ Software (images acquired at 400 ms intervals). Acquisition parameters were controlled using a Slidebook 6.0 program (3i - Intelligent Imaging).

### FRAP

The FRAP experiment was conducted using a Yokogawa CSU-W1 SoRa dual cam spinning disk confocal. The microscope is equipped with an Acal BFi UV Opti-Microscan point scanner that we used for the FRAP experiment. This system is integrated with Nikon NIS Elements software for seamless experimental setup and data acquisition. The movies were acquired with a 60 × 1.27 NA water objective and 2.8× SoRa magnifier with 100 ms exposure times and 250 ms intervals (4 frames/s).

FRAP was calculated with a combination of Fiji (*76*) and MATLAB (MATLAB and Statistics Toolbox Release 2012, The MathWorks, Nitick, USA). Time-lapse images of spindles expressing GFP:: α-tubulin and mCherry::histone (corresponding to chromosomes) were realigned in a routine for matching, rotation, and translation using Rigid Body of Fiji’s StackReg plug-in, so that the random displacement of the spindle due to the spontaneous motion of the worm was corrected.

### Sample preparation for electron tomography

Wild-type (N2) *C. elegans* embryos were collected in cellulose capillary tubes (*77*) and high-pressure frozen as described using an EM ICE high-pressure freezer (Leica Microsystems, Vienna, Austria) (*78*). Freeze substitution was performed over 2–3 d at –90°C in anhydrous acetone containing 1% OsO_4_ and 0.1% uranyl acetate. Samples were embedded in Epon/Araldite and polymerized for 2 d at 60°C.

For control 1 and 2 spindles, serial semi-thick sections of 300nm were cut using an Ultracut UCT Microtome (Leica Microsystems, Vienna, Austria) collected on Formvar-coated copper slot grids and post-stained with 2% uranyl acetate in 70% methanol followed by Reynold’s lead citrate. For dual axis tomography (*79*) 15 nm colloidal gold particles (Sigma-Aldrich) were attached to both sides of semi-thick sections to serve as fiducial markers for subsequent image alignment. Series of tilted views were recorded using a Tecnai F30 transmission elctron microscope (FEI Company, Eindhoven, The Netherlands) operated at 300 kV and equipped with a Gatan CCD camera (2k x 2k). Images were captured every 1.0° over a ±60° range.

For control 3 and *klp-19 (RNAi)* 1 and *klp-19 (RNAi) 2,* Serial semi thick sections (200 nm) were cut using a Leica Ultracut S microtome (Leica Microsystems, Vienna, Austria) collected on Pioloform-coated copper slot grids and post stained with 2% uranyl acetate in 70% methanol followed by Reynold’s lead citrate. For dual-axis electron tomography, 15 nm colloidal gold particles (Sigma-Aldrich) were attached to both sides of semi-thick sections to serve as fiducial markers for subsequent image alignment. A series of tilted views were recorded using an F20 electron microscopy (Thermo-Fisher, formerly FEI) operating at 200 kV using a Gatan US4000 (4000 px × 4000 px) CCD (for control 3) or a Tietz TVIPS XF416 camera (for *klp-19 (RNAi)* 1 and 2. Images were captured every 1.0° over a ±60° range.

### Quantification of electron tomography data

We used the IMOD software package (http://bio3d.colorado.edu/imod) for the calculation of electron tomograms (*80*). We applied the Amira software package for the segmentation and automatic tracing of microtubules (*81*). For this, we used an extension to the filament editor of the Amira visualization and data analysis software (*82, 83*). We also used the Amira software to stitch the obtained 3D models in *z* to create full volumes of the recorded spindles (*84*). The automatic segmentation of the spindle microtubules was followed by a visual inspection of the traced microtubules within the tomograms. Correction of the individual microtubule tracings included manual tracing of undetected microtubules, connection of microtubules from section to section, and deletions of tracing artifacts (e.g., membranes of vesicles).

### Data analysis for tomography

Data analysis was performed using the Amira software (Visualization Sciences Group, Bordeaux, France).

#### Length distribution of microtubules

For the analysis of the microtubules length distributions, we previously found that removing MTs that leave the tomographic volume only had minor effects on the length distribution (*47*). Therefore, we quantified all microtubules contained in the volume. In addition, in all analyses, microtubules shorter than 100 nm were excluded to reduce errors due to the minimal tracing length.

#### Interaction analysis

For the detection of possible interactions in 3D, the Interaction and Distance modules of the SpindleAnalysis Toolbox were used in Amira.

Interactions computes possible microtubule intersections. An intersection is defined as a part of a microtubule that is closer to another microtubule than given a threshold (here 100nm). In detail, for each point of a microtubule, the distance to another microtubule is computed. Based on these distances, all parts of the microtubule are detected that lie closer to the other microtubule than the threshold. For each part an interaction is generated and stored in a spread sheet. For each interaction the approximated angle and the interaction start, end and length is scored, In addition, we defined a minimal interaction length of 100nm, over which the distance between the interacting microtubules has to remain within the defined threshold distance for an interaction (here 100nm) to filter the interactions.

The distance module computes the distance between microtubules. For each point of a microtubule the closest segment of all other microtubules is detected and stored as a floating point attribute. Furthermore, for each microtubule, the closest other microtubule is detected. The distance between two microtubules is defined as the minimal distance between the segments. The ID and the distance are stored as attributes on the edges.

For each line segment of a microtubule the distance to the surrounding microtubules was computed analytically. The distance of a microtubule was defined as the minimum of all segment distances. Second, for each pair of microtubules the distance and the angle were computed. The distance between two microtubules was defined as the minimum of the distances between all their line segments. A 3D grid data structure was used to accelerate these computations. To reduce errors due to local distortions of the microtubules, the angle is defined by the angle between the lines through the start and end points of the microtubules. Third, based on these data an abstract graph was constructed, where each microtubule is represented as a vertex and each interaction (based on thresholds for interaction distance and angle) as an edge.

### Quantitative analysis of Microtubule growth in *C. elegans* mitotic spindles

EB-2 GFP normalized mean intensity was measured in a fixed-size ROI in the midzone region. EB-2 GFP binding events were counted in a similar way to Srayko et al., 2005 (*85*). Kymographs were generated in Fiji based on the lines half-distance between anterior centrosome and cortex.

Segment of 4.5µm and 10% thresholding was used to segment EB-2 binding events and count them. Spd1-GFP and EB2-GFP profiles were measured with a mean profile intensity of the ROI box across the spindle.

### Statistical analysis

Statistics are presented as mean ± SEM or mean ± STD, and *p* values were calculated by the “ttest2” function in MATLAB.

### Second harmonic generation imaging and two-photon florescence imaging

Simultaneous SHG imaging and TP imaging were constructed around an inverted microscope (Eclipse Ti, Nikon, Tokyo, Japan), with a Ti:sapphire pulsed laser (Mai-Tai, Spectra-Physics, Mountain View, CA) for excitation (850 nm wavelength, 80 MHz repetition rate, ∼70 fs pulse width), a commercial scanning system (DCS-120, Becker & Hickl, Berlin, Germany), and hybrid detectors (HPM-100-40, Becker & Hickl). The maximum scan rate of the DCS-120 is ∼2 frames/s for a 512 × 512 image. The excitation laser was collimated by a telescope to avoid power loss at the XY galvanometric mirror scanner and to fill the back aperture of a water-immersion objective (CFI Apo 40× WI, NA 1.25, Nikon). A half-wave plate (AHWP05M-980) and a quarter-wave plate (AQWP05M-980) were used in combination to achieve circular polarization at the focal plane, resulting in equal SHG of all orientations of microtubules in the plane, unbiased by the global rotation of the spindle, the spatial variation in the angle of the microtubules, and the local angular disorder of microtubules. Forward-propagating SHG was collected through an oil-immersion condenser (1.4, Nikon) with a 425/30 nm filter (FF01-425/30-25, Semrock). Two-photon fluorescence was imaged with a non-descanned detection scheme with an emission filter appropriate for green-fluorescent-protein (GFP)-labeled tubulin in *C. elegans* (FF01-520/5-25, Semrock, Rochester, NY).

Both pathways contained short-pass filters (FF01-650/SP-25, Semrock) to block the fundamental laser wavelength. Image analysis was performed with MATLAB (The MathWorks, Natick MA), and ImageJ (National Institutes of Health, Bethesda, MD.

### Mitotracker

Timelapse fluorescent image sequences were processed in Python using nd2reader (https://github.com/Open-Science-Tools/nd2reader), OpenCV (https://pypi.org/project/opencv-python/), and scipy (https://scipy.org) along with other standard packages for data analysis.

Multi-channel images were converted to 8-bit per channel and then split by spindle and DNA channels. Embryos remained in a relative fixed position during observation. Outlines of individual embryos were determined by creating a median projection of low background fluorescence of the GFP microtubule signal along the time axis, followed by watershed segmentation. The embryo outlines were used to create masks to eliminate all fluorescent microtubule and chromatid signals external to the individual embryo. To identify the absolute position of spindle poles and chromatids, a gaussian blur with a 3x3 kernel was applied to each channel at each timepoint, followed by binary thresholding using the top 25% (191–255) and top 50% (127–255) signal intensity for spindle poles and chromatid, respectively. Positions of each spindle poles and chromatid groups were determined as center or edge of the binarized objects, respectively. The midzone was determined as the half-distance between both spindle poles. The distance and velocity of the edge of the leading chromatids (advancing toward each spindle pole) was plotted over time relative to this midzone position. The oscillation of each spindle pole was tracked and plotted independently relative to a fixed reference axis. By default, this reference axis corresponded to the long axis of the embryo; however, the axis could be adjusted to align with the spindle axis of any chosen timepoint.

Kymographs for spindle pole and chromatid movements were created by applying a rotation to each frame such that the spindle pole-to-spindle pole axis in each frame aligns with a horizontal reference line. Each line in the final kymograph reflects a sum intensity projection of 10 pixels along the y-axis (+/- 5 pixel above/below the reference axis) of each rotated image. Enhanced kymographs were created from the original kymographs by applying standard adaptive thresholding and skeletonization to visualize the trajectories of spindle pole centers and chromatids’ leading edges. The Python code is available here: https://github.com/uvarc/mitosisanalyzer. Tracked data was used to plot distances between centrosomes and chromosomes at a specific time or during anaphase. Segregation rates were calculated from the displacement during first 40s after the beginning of anaphase.

### Microfluidic device fabrication

The microfluidics were fabricated using an SU-8 photolithography similar to previous studies (*62, 64, 86*). Briefly, a 10:1 mixture was made using PDMS and a curing agent, mixed and poured in a mold. The molds were degassed for 2min, spin coated with PDMS at 300 rpm for 30 seconds and baked at 85℃ for 20 mins. Solidified PDMS was then peeled off and punched with a 2 mm biopsy punch to make inlet and outlet wells.

A 24 x 60 mm glass coverslip was cleaned with compressed nitrogen gas to remove visible dust and placed in a UV-ozone for 20 min for further cleaning. PDMS was also plasma treated for 30 seconds and immediately placed on the top of UV-ozone treated coverslip to create a permanent bond. The devices were then immediately passivated by adding 2 μL of PEG solution (monomethyl -PEG-silane with 2% HPLC water and 0.02% biotin-PEG-silane) and incubated at room temperature for 60 mins. After incubation, excess PEG was aspirated using a vacuum, rinsed 2x with water and dried in an airtight container with desiccant for at least 24 hrs before use.

For experiments in Figure 4B and Figure 4A-C, the devices were functionalized with *in vitro* polymerized microtubules immediately before use. First, the device was rehydrated by flowing through 2 μL of water and aspirating it using a vacuum from the outlet while leaving some in the channel. Second, a solution of 0.2 mg/mL of neutravidin in water was incubated in the device for 10 min. Post incubation the device was rinsed 3x with water and 1x with tubulin buffer (80 mM Pipes pH 6.9, 1 mM EGTA, 1 mM MgCl2, 10% glycerol) plus 20 μM Paclitaxel (SIGMA 33069-62-4). Third, 2 μL of polymerized microtubule solution were added in the inlet and immediately aspirated out from the outlet to prevent creation of a mesh of microtubules at the inlet that blocks flow. An additional 2μL of microtubule solution was added to the inlet and incubated at room temperature for 15 mins. Post incubation, the device was rinsed 4x with tubulin buffer containing 20 μM pacitaxel. After rinsing, 0.5 μL of tubulin buffer was added to the inlet and outlet wells and the device was sealed with clear tape. Before use, the tape was cut off using a scalpel and peeled using forceps.

For experiments in Figure S8C and Figure 4C, the device was functionalized following the sc-SiMPull standard antibody functionalization (*62*). Immediately before use the device was rehydrated using SiMPull Buffer (10 mM Tris (pH 8.0), 50 mM NaCl, 0.1% Triton X-100, and 0.1 mg/mL bovine serum albumin). A 0.2 mg/mL solution of neutravidin was added to each well and incubated at room temperature for 10 mins. The device was then rinsed 4x with SiMPull Buffer. After the washing step, a 100 μM of anti-GFP antibody was added to each well and incubated at room temperature for 10 mins. The device was rinsed 4x to remove excess antibodies. The channels that were not immediately used were hydrated with 0.5 μL of SiMPull buffer and sealed with clear tape to prevent evaporation.

### Tubulin Polymerization

Stock tubulin was made by mixing cycled-tubulin, Alexa 647-tubulin and biotin-tubulin (Pursolutions: 032005, 064705, 03305, respectively) on ice to a final concentration of 20 mg/mL: 2 mg/mL: 0.8 mg/mL, respectively. This mixture was spun down in an ultracentrifuge at 70,000 rpm at 4℃ for 7 min to remove aggregated protein. The supernatant was removed and aliquoted in 2 μL volume, snap frozen in liquid nitrogen and stored at -80℃.

To polymerize tubulin, 2 μL of the stock tubulin was added to 6 μL of tubulin buffer containing 1 mM GTP (final concentration of 5 mg/mL cycled tubulin; 0.5 mg/mL Alexa 647-tubulin and 0.2 mg/mL of Biotin-tubulin). This mixture was first incubated on ice for 5 min to promote nucleotide binding, then incubated in a 37°C water bath for 45 mins. After polymerization, microtubules were stabilized by adding paclitaxel at a final concentration of 35 μM and incubated for 20 mins at 37℃. After paclitaxel stabilization, this stock of microtubule was diluted at 1:100 in tubulin buffer containing 20 μM paclitaxel and used to functionalize the microfluidic device. Post polymerization all tubulin buffer must be supplemented with 20 μM paclitaxel.

### sc-SiMPull and TIRF microscopy

Gravid adults were dissected in 1x egg buffer ((5 mM HEPES pH 7.4, 118 mM NaCl, 40 mM KCl, 3.4 mM MgCl2, 3.4 mM CaCl2) and embryos were rinsed 2x in tubulin buffer containing 20 μM paclitaxel and 500 μM ATP. Embryos were rinsed by mouth pipetting them into depression slides containing ∼ 30 μL of the aforementioned solution. In parallel, the microfluidic device was rinsed 2x with tubulin buffer containing paclitaxel and ATP, and an embryo was placed in the inlet. The embryo was pulled into the channel by aspirating the outlet using a vacuum. Both the inlet and the outlet were sealed with small pieces of clear tape to prevent fluid evaporation and flow. The device was then transferred to a TIRF microscope for laser lysis and imaging.

For experiments in Figure S4C-D, SiMPull buffer was used instead of tubulin buffer.

TIRF images in Figure 4B, Figure S8C, and Figure 4A were acquired using a Nikon Eclipse Ti-2 microscope equipped with 100X 1.49 NA objective and a Photometrics Prime 95B camera. TIRF images in Figure 2B were acquired on 60X 1.49 NA objective and a Photometrics Prime BSI camera. TIRF illumination was achieved by an iLas2 circular TIRF illuminator (Roper scientific, E’vry France). To this microscope, we added an infrared pulsed laser (STP-40K, Teem photonics, Grenoble, France; Arduino, Somerville, MA) that was installed as a separate path and focused to the sample. GFP was excited using a 488 nm laser, and Alexa 647 was excited using a 638 nm laser.

TIRF images in Figure 4C were acquired using a home-built micromirror TIRF microscope based on the MadCity Labs RM21 platform. Excitation light from 488 nm and 638 nm lasers was collimated, combined, and focused onto a 2 mm-diameter micromirror that coupled the excitation light to the back aperture of a 60x, 1.50 NA TIRF objective lens (Olympus UPLAPO60XOHR). A second micromirror, positioned on the opposite side of the objective, collected the totally internally reflected excitation light. The emitted light passed between the two micromirrors and was passed through custom multicolor image splitting optics before being captured on a Photometrics PrimeBSI Express camera.

### Image processing and particle tracking

FIJI was used to adjust the brightness and contrast for visibility, to make kymographs, and maximum time projections.

Single particles were tracked using Utrack (*87*) and modified the following parameters: point detection with alpha value of 0.05, Brownian+ directed motion mode with minimum track length of 2 and maximum GAP to close of 2. All other parameters were left as default. Detailed protocol was published in (*88*). Data from Utrack data was then analyzed using homemade MATLAB scripts to determine the velocity, brightness and run length sc-SiMPull data were analyzed using the SiMPull analysis MATLAB package (Dickinson et al. 2017; Sarikaya et al. 2021) which is freely available at https://github.com/dickinson-lab/SiMPull-Analysis-Software. Briefly, the software detects each single-molecule capture event in a sc-SiMPull dataset; determines the stoichiometry of the bait protein by counting photobleaching steps; and checks whether or not the molecule co-appears with prey protein signal.

## Supplementary Figure Legends

**Supplementary Figure 1.**
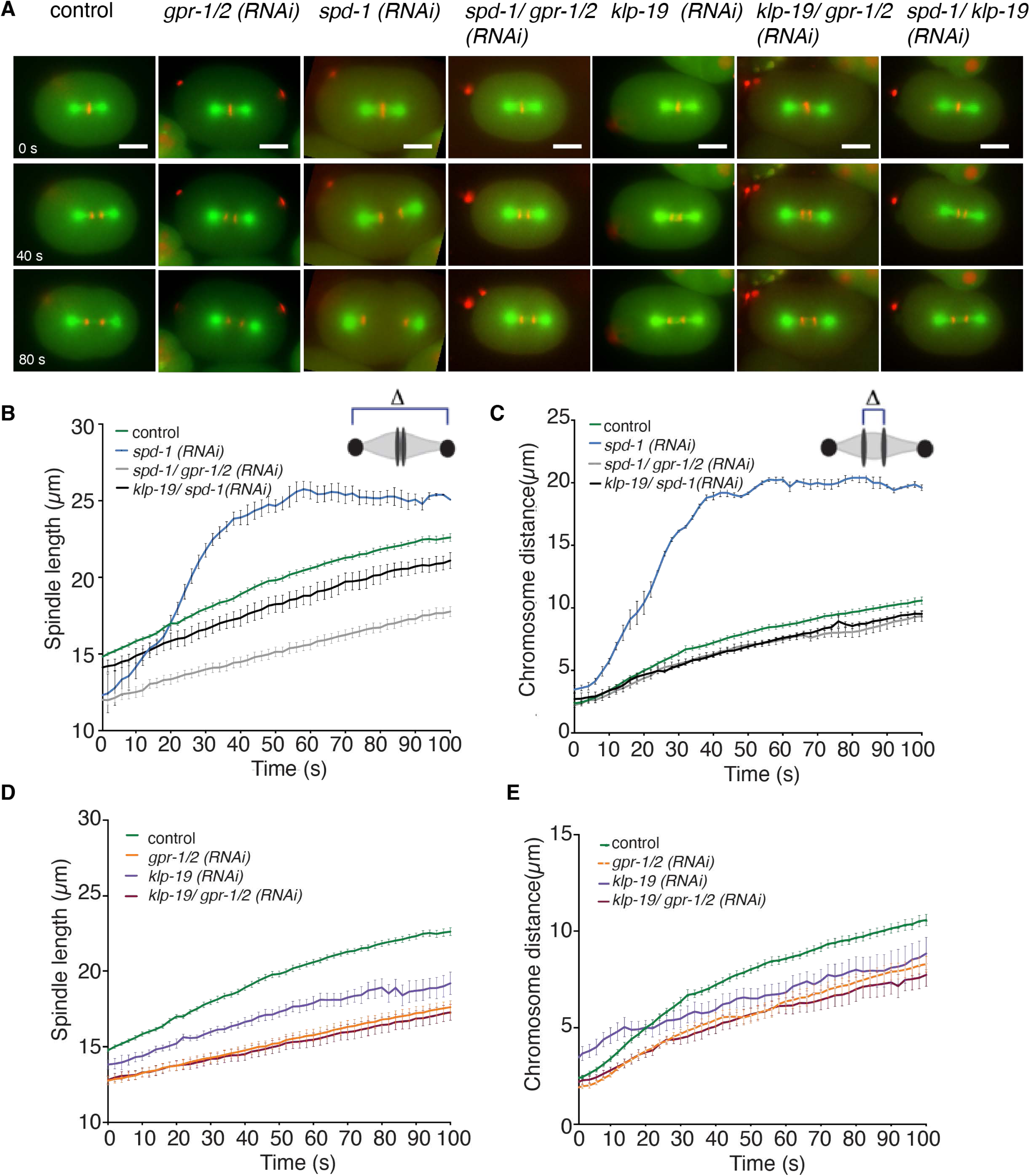
Depletion of midzone components affects spindle dynamics. **A.** Stills of embryos in control embryos and after different RNAi treatments at 0s, 40s and 80s after anaphase onset. Embryos are expressing α-tubulin::GFP and histone::mCherry. Scale bar is 10µm. **B.** Plot of pole-to-pole distance throughout anaphase in control embryos (green), after *spd-1 (RNAi)* (blue), *klp-19/ spd-1 (RNAi)* (black) and *spd-1/ gpr-1/2 (RNAi)* (grey). Timepoint “0” is defined as anaphase onset **C.** Plot of chromosome distance throughout anaphase in control embryos (green), after *spd-1 (RNAi)* (blue), *klp-19/ spd-1 (RNAi)* (black) and *spd-1/ gpr-1/2 (RNAi)* (grey). **D.** Plot of pole-to-pole distance throughout anaphase in control embryos (green), after *klp-19 (RNAi)* (purple), *gpr-1/2 (RNAi)* (orange) and *klp-19/ gpr-1/2 (RNAi)* (maroon). **E.** Plot of chromosome distance throughout anaphase in control embryos (green), after *klp-19 (RNAi)* (purple), *gpr-1/2 (RNAi)* (orange) and *klp-19/ gpr-1/2 (RNAi)* (maroon). Error Bars are sem.

**Supplementary Figure 2.**
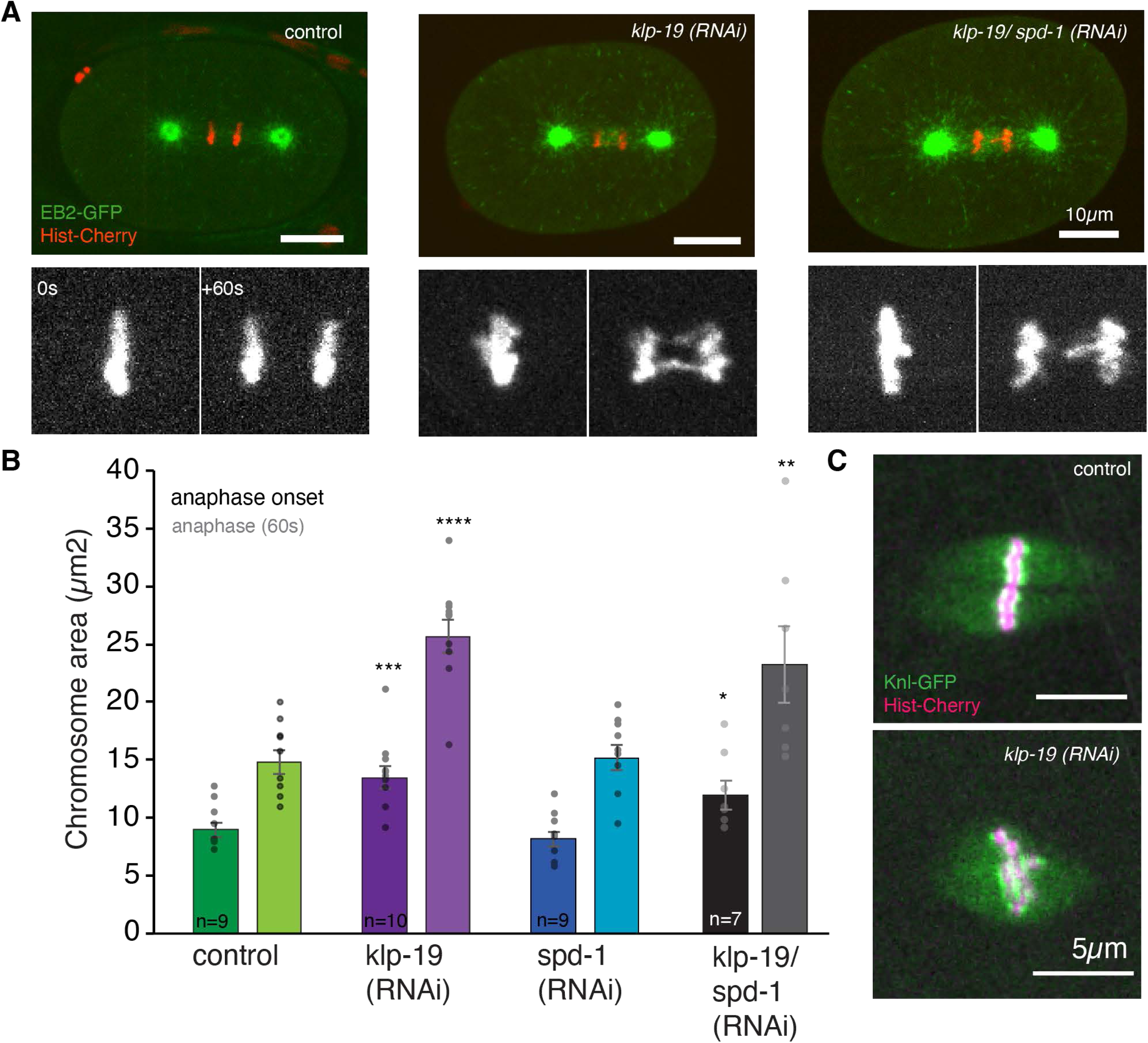
Depletion of KLP-19 affects chromosome segregation. **A.** Stills of control embryos and after different RNAi treatments at 60s after anaphase onset. Embryos are expressing EBP2::GFP (green) and histone::mCherry (red). Bottom images show chromosomes at anaphase onset and 60s after anaphase onset **B.** Bar plot of the chromosome area at metaphase and 60s after anaphase onset in control embryos and after different RNAi treatments. **C**. Stills of control embryos and embryos after *klp-19 (RNAi)* expressing kinetochore marker KNL-1::GFP and mCherry histone . Scale bar is 10µm in A and 5µm in C. Error Bars are sem. The significance of differences between control and RNAi conditions was determined by two-tailed Student’s *t*-tests (\*\*\*\**P* < 0.0001).

**Supplementary Figure 3.**
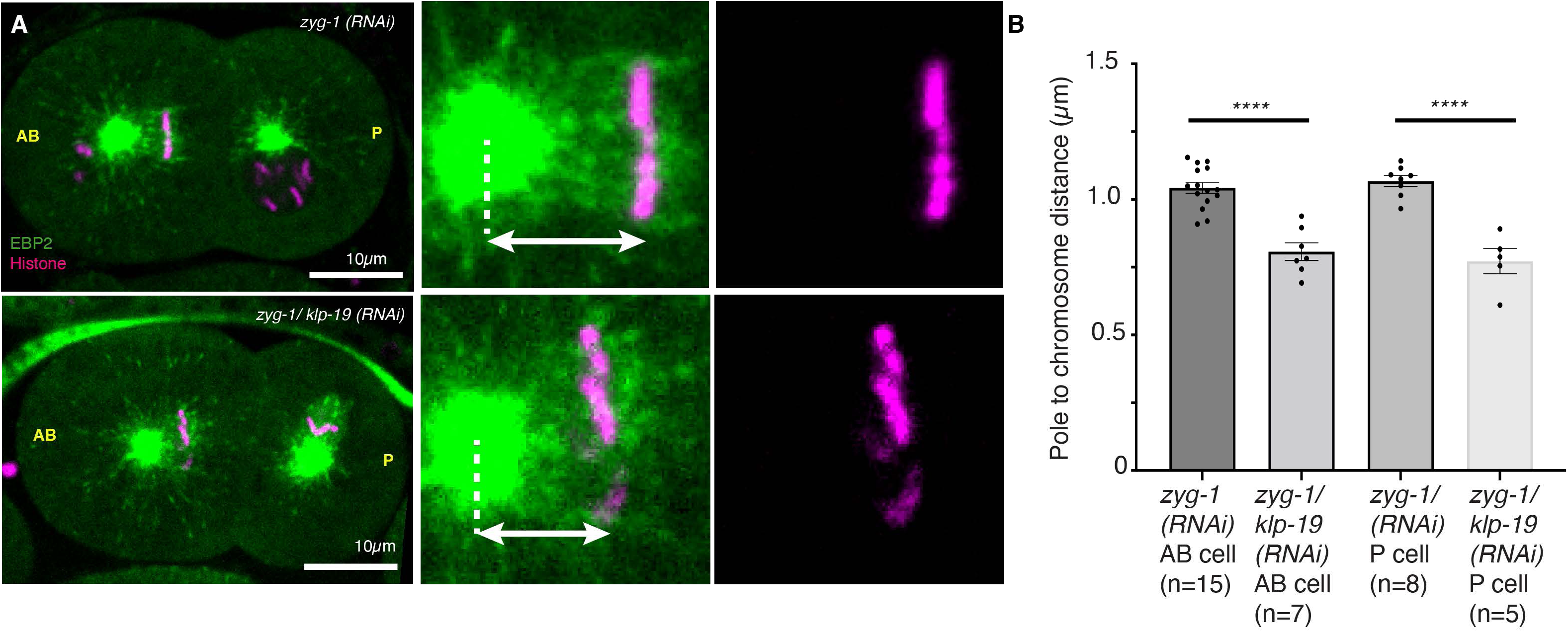
KLP-19 affects polar ejection forces. **A.** Stills of embryos treated with *zyg-1 (RNAi)* to induce the formation of monopolar spindles in the 2-cell stages. Embryos are expressing EBP-2::GFP (green) and mCherry::Histone. (magenta) Top shows *zyg-1 (RNAi)* embryos, bottom *zyg-1/ klp-19 (RNAi)* treated embryos. Scale bar 10µm **B**. Plot of the distance between centrosomes and metaphase plate in each of the 2-cells (AB and P) in control and *klp-19 (RNAi)* embryos. Error Bars are sem. The significance of differences between control and RNAi conditions was determined by two-tailed Student’s *t*-tests (\*\*\*\**P* < 0.0001).

**Supplementary Figure 4.**
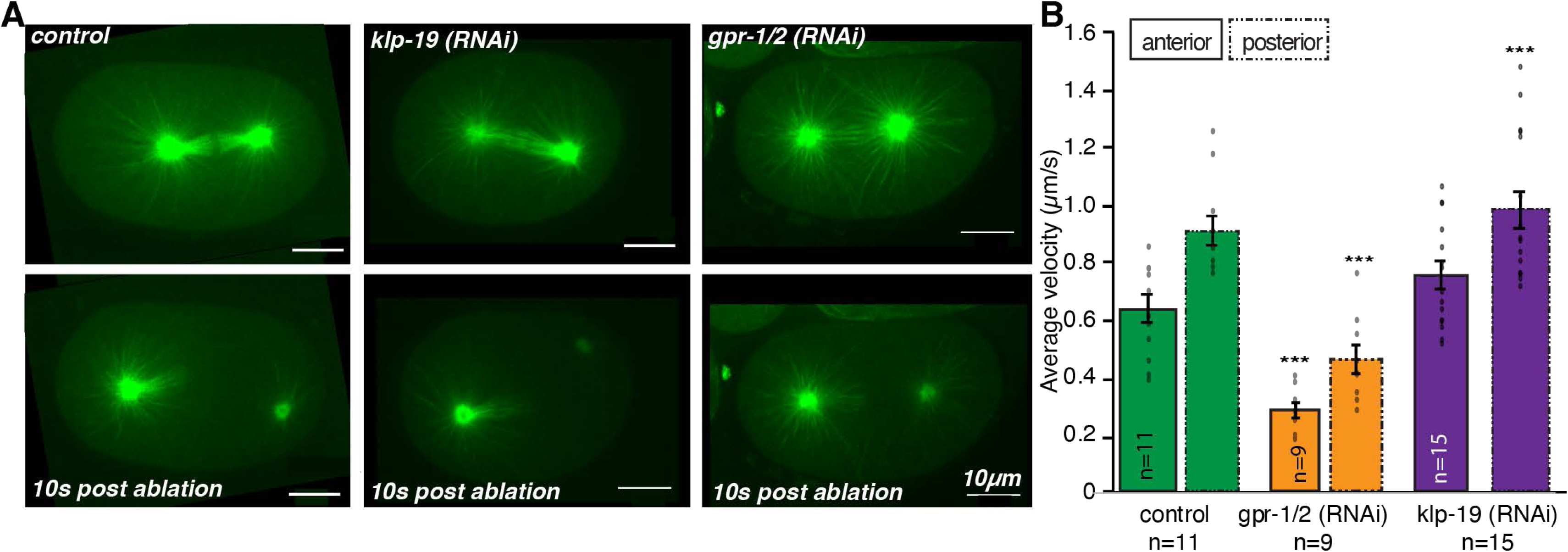
KLP-19 depletion does not directly affect cortical pulling forces. **A.** Stills of embryos before (top) and 10 s after (bottom) laser microsurgery severing the posterior centrosome from the spindle in control (left), *klp-19 (RNAi)* (middle) and *gpr-1/2 (RNAi)* (right) treated embryos. **B.** Plot of the velocity of the anterior (continuous line) and posterior (dashed line) centrosome after laser microsurgery in control (green), *gpr-1/2 (RNAi)* (orange) and *klp-19 (RNAi)* (purple) treated embryos. The significance of differences between control and RNAi conditions was determined by two-tailed Student’s *t*-tests (\*\*\*\**P* < 0.0001). Error bars are sem, Scale bar is 10µm.

**Supplementary Figure 5.**
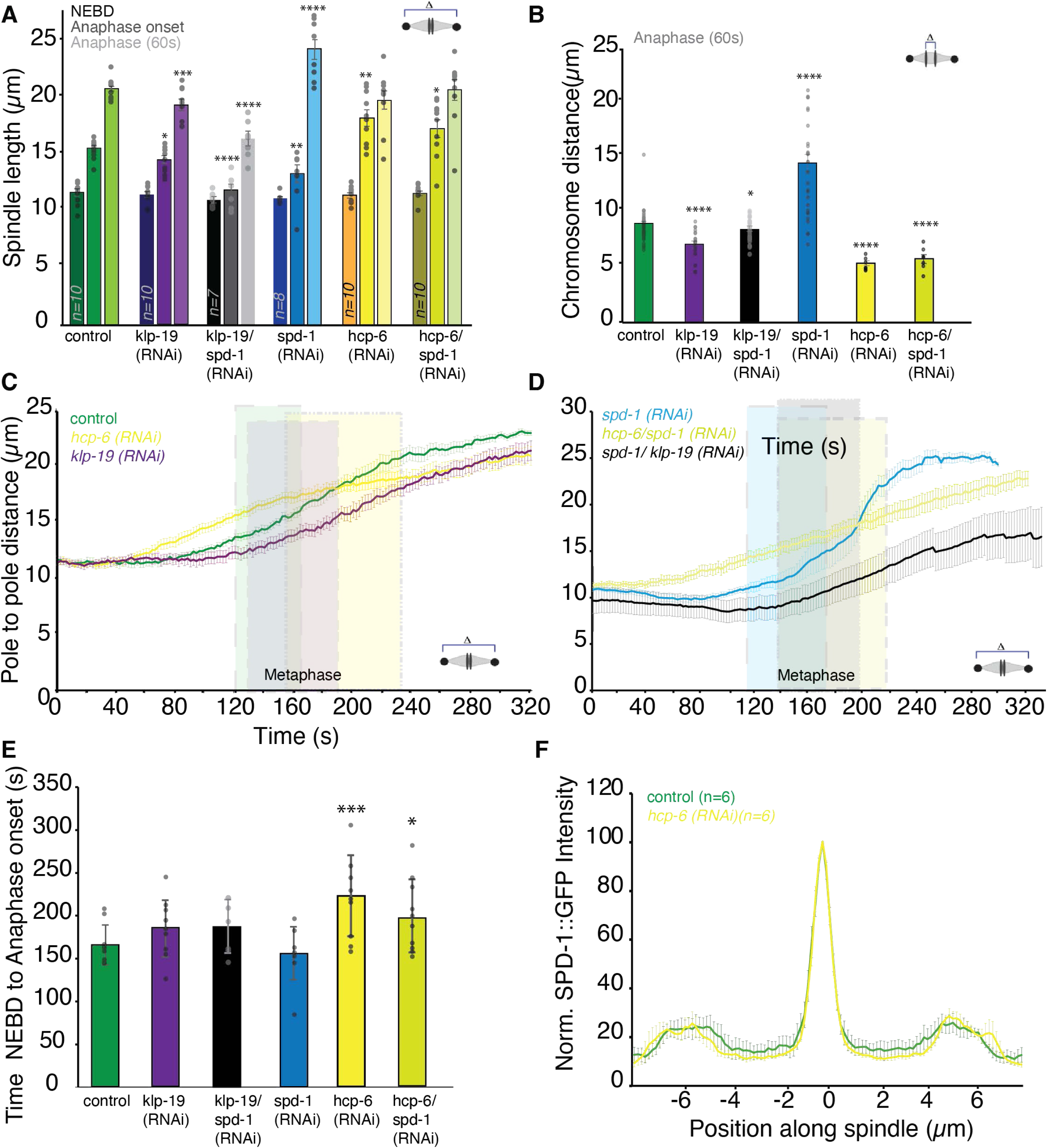
Spindle dynamics in *hcp-6 (RNAi)* treated embryos. **A.** Plot of the spindle length at NEBD, Anaphase onset and 60s after anaphase onset in control embryos and embryos depleted of KLP-19, KLP-19/ SPD-1, SPD-1, HCP-6 and HCP-6/ SPD-1. **B.** Plot of the chromosome distance at 60s after anaphase onset. **C.** Plot of the spindle length from NEBD (0) through metaphase and anaphase in control *hcp-6 (RNAi)* and *klp-19 (RNAi)* treated embryos. The average timepoint of metaphase is indicated by the shaded boxes. **D.** Plot of the spindle length from NEBD (0) through metaphase and anaphase in *spd-1 (RNAi), hcp-6/ spd-1 (RNAi)* and *spd-1/ klp-19 (RNAi)* treated embryos. The average timepoint of metaphase is indicated by the shaded boxes. **E.** Bar plot of the average time from NEBD to anaphase onset for control and RNAi treated embryos. **F.** Plot of SPD-1 GFP intensity along the spindle axis in control embryos (green) and embryos after HCP-6 depletion (yellow). Color code: control embryos = green, *klp-19 (RNAi)* = purple, *klp-19/ spd-1 (RNAi) =* grey, *spd-1 (RNAi)* = blue, *hcp-6 (RNAi)* = yellow, *hcp-6/ spd-1 (RNAi)* = light green. The significance of differences between control and RNAi conditions was determined by two-tailed Student’s *t*-tests (\*\*\*\**P* < 0.0001). Error bars are sem.

**Supplementary Figure 6.**
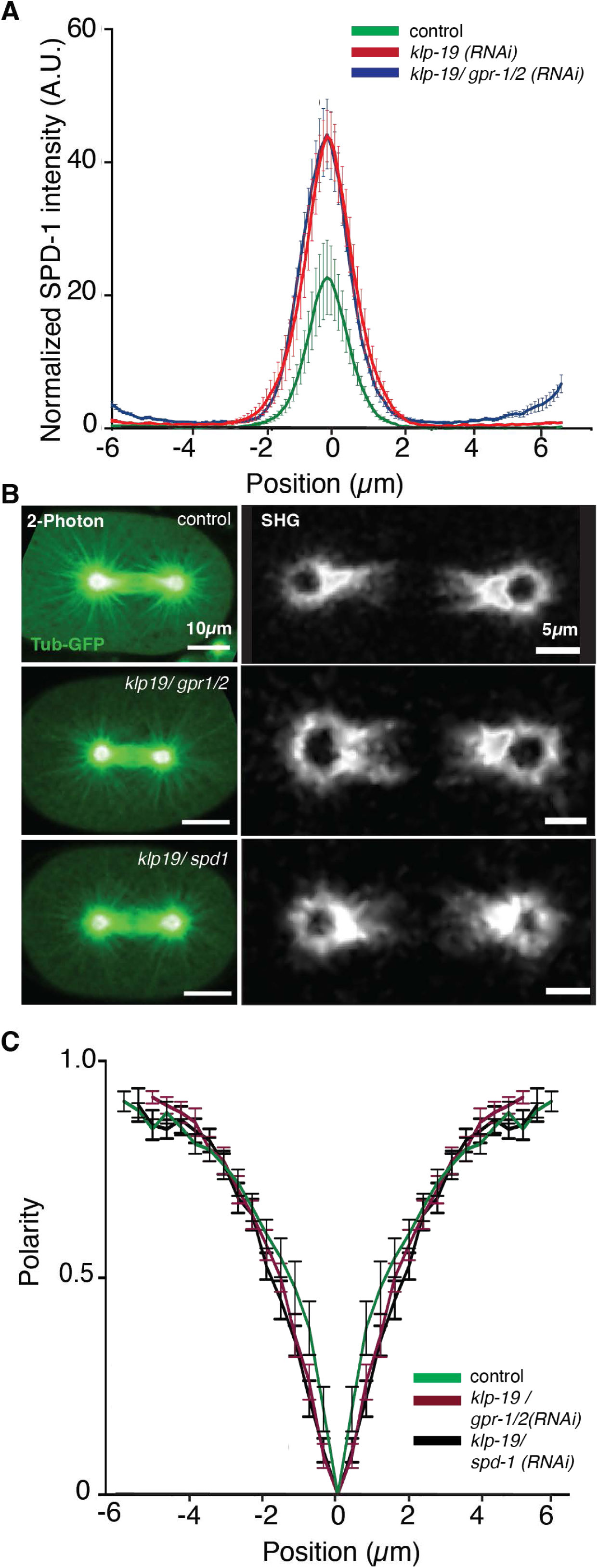
KLP-19 affects microtubule overlap in the spindle midzone. **A.** Plot showing the normalized intensity of SPD-1 GFP along the spindle axis in control embryos (green), embryos depleted of KLP-19 (red) and embryos after *klp-19/ gpr-1/2 (RNAi)* (blue). “0” is the spindle center **B.** Left: Two-photon microscopy images of α-tubulin::GFP in control (top), *klp-19/ gpr-1/2 (RNAi)* and klp-19/ spd-1 (RNAi) treated embryos. Right: Corresponding SHG images. Scale Bars are 10µm left, 5µm right images **C.** Plot of the polarity of microtubules along the spindle axis (0= spindle center) of control embryos and after *klp-19/ gpr-1/2 (RNAi)* and *klp-19/ spd-1 (RNAi).* Error bars are sem.

**Supplementary Figure 7.**
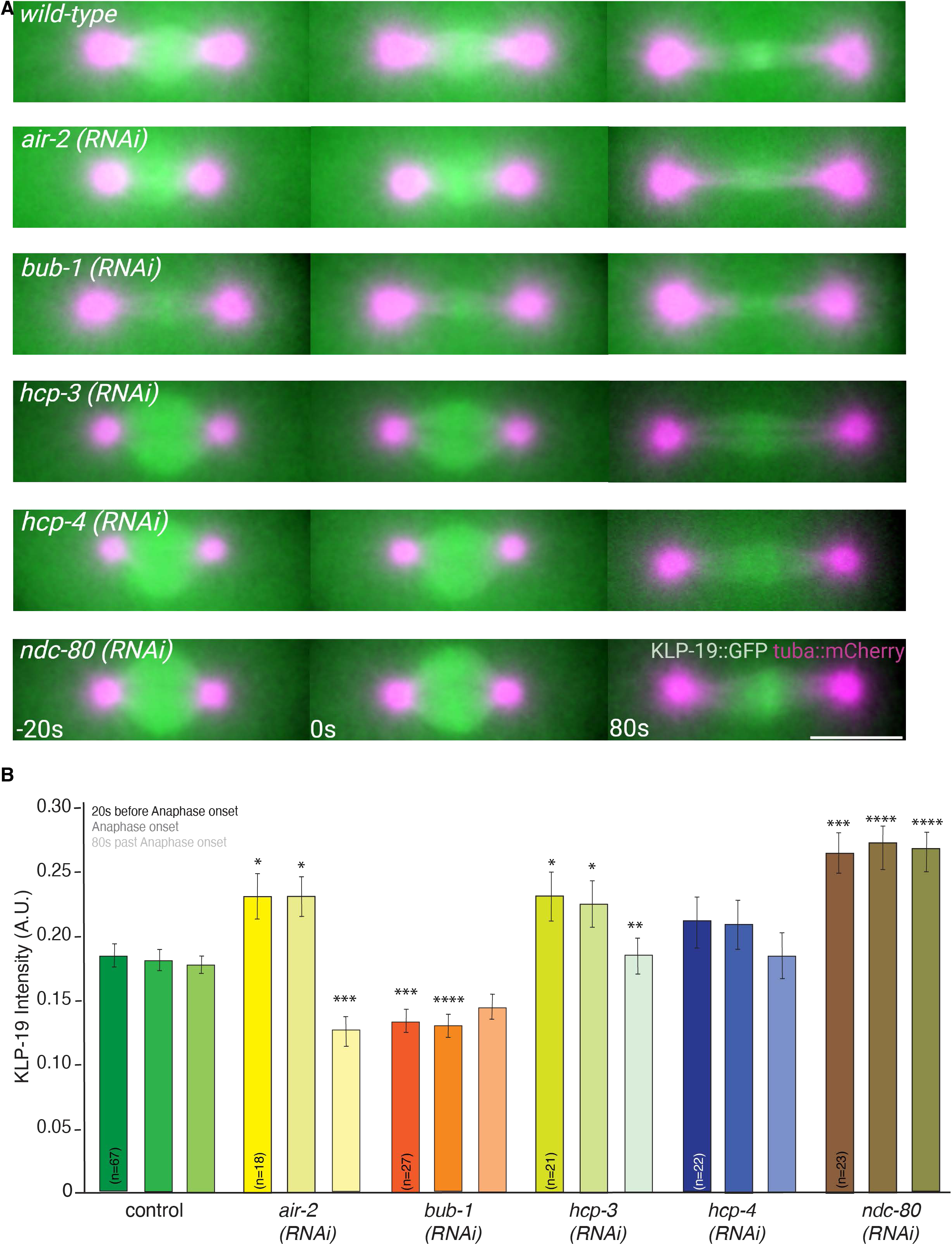
KLP-19 localization to the spindle midzone depends on BUB-1 and AIR-2. **A.** Stills of control embryos (top), embryos treated with different RNAis throughout anaphase in embryos expressing α-tubulin::mCherry and KLP-19::GFP. **B.** Bar plot of the normalized KLP-19 intensity in the spindle center at 20s before metaphase, metaphase and 80s after anaphase onset. Color code: control embryos = green, *air-2 (RNAi)* = yellow, *bub-1 (RNAi) =* red, *hcp-3 (RNAi)* = light green, *hcp-4 (RNAi)* = blue, *ndc-80 (RNAi)* = brown. The significance of differences between control and RNAi conditions was determined by two-tailed Student’s *t*-tests (\*\*\*\**P* < 0.0001). Error bars are sem.

**Supplementary Figure 8.**
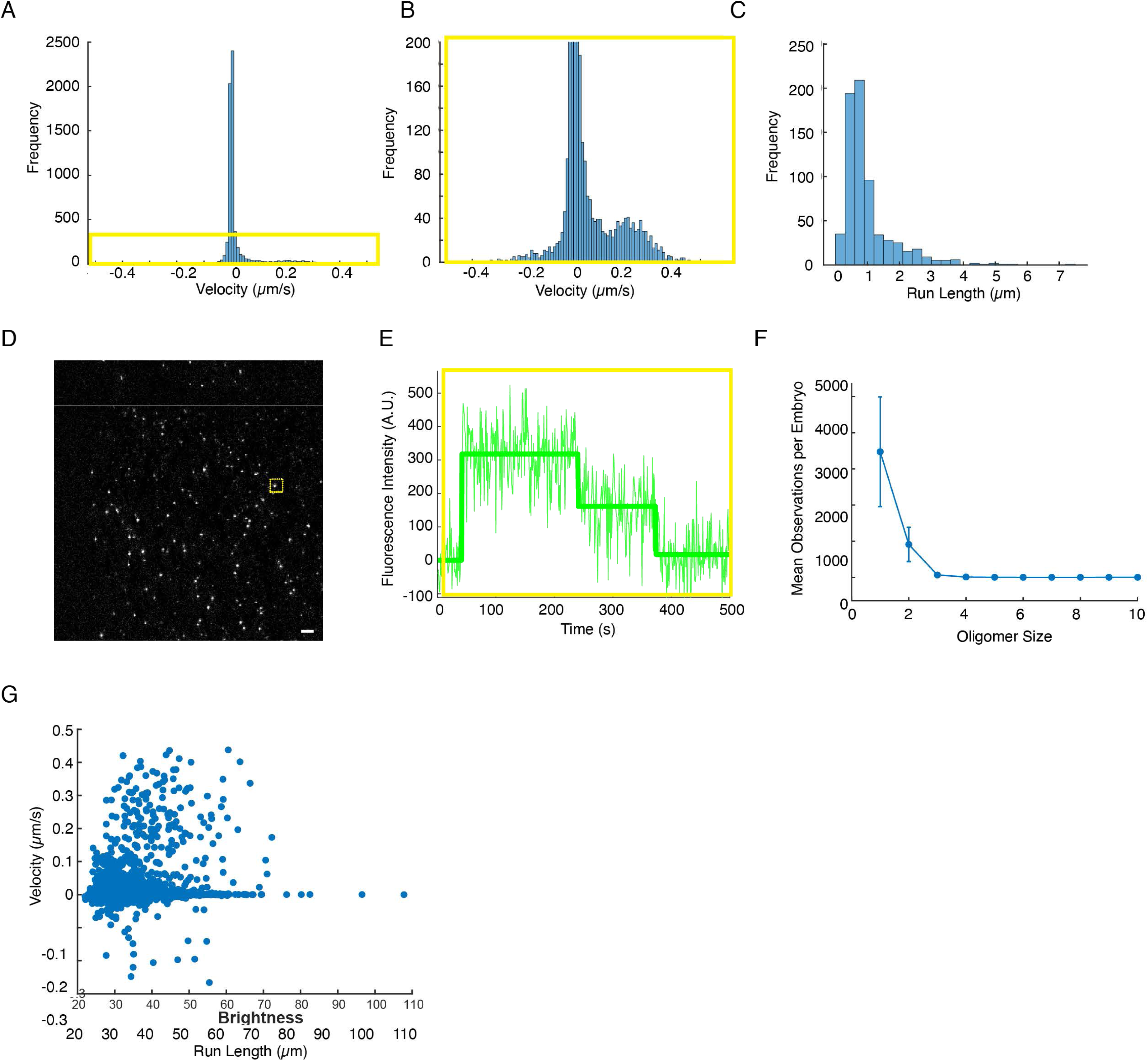
KLP-19 molecules form homodimers. **A**. Histogram plot showing the velocity of KLP-19. **B.** Enlarged region highlighted from the left panel in yellow. **C** Histogram plot showing KLP-19 run length on microtubules. **D**. TIRF difference image showing newly appeared KLP-19 molecules. **E**. Example intensity trace of the single particle highlighted in yellow, showing two photobleaching steps. **F**. Plot showing the frequency of different KLP-19 oligomer sizes. **G**. Plot showing KLP-19 velocity as a function of particle brightness.

**Supplementary Figure 9.**
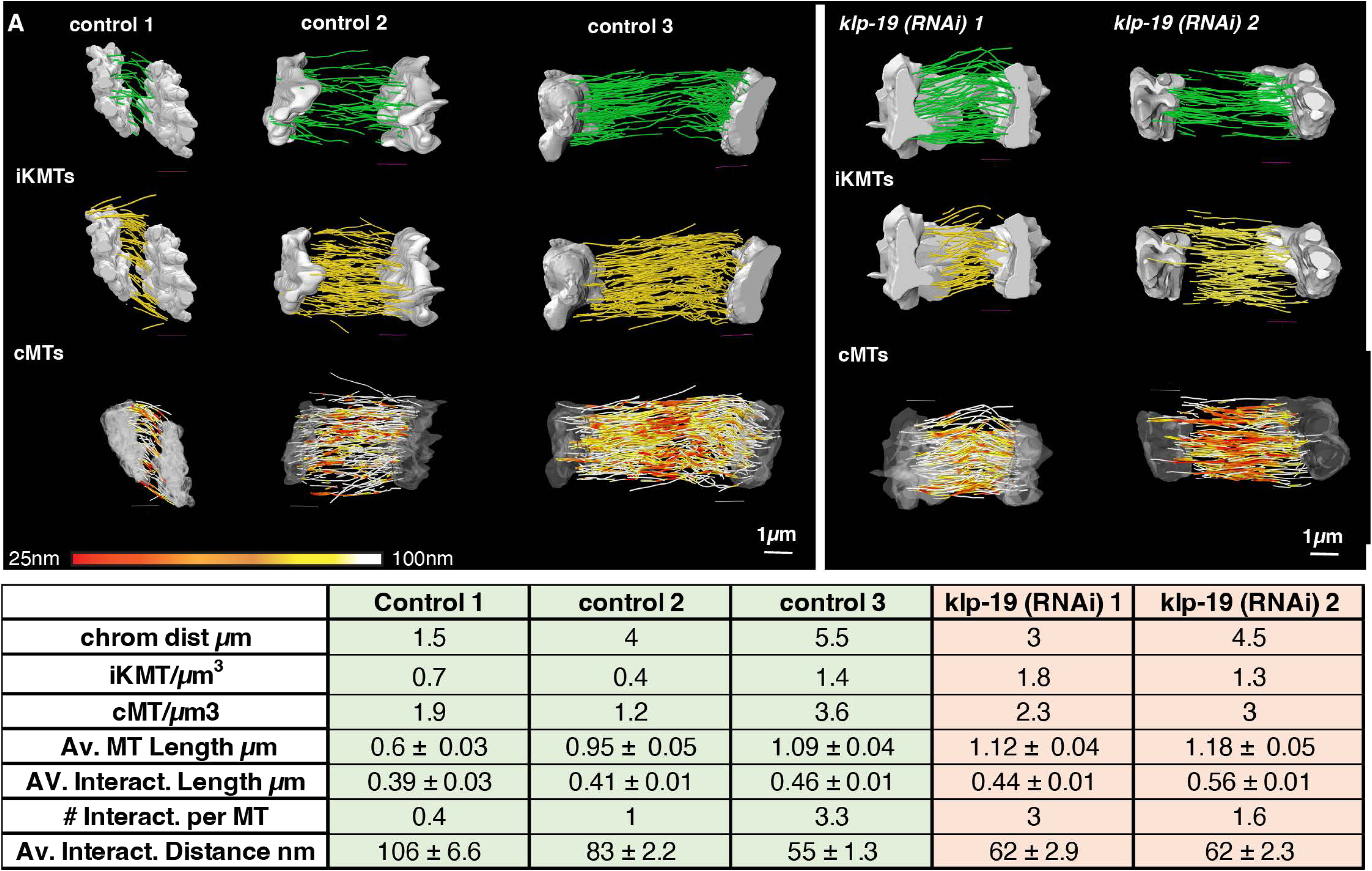
KLP-19 depletion leads to increased microtubule length and interactions. **A.** 3D tomographic reconstruction obtained by electron tomography of all control and *klp-19 (RNAi)* embryos showing iKMTs (top), cMTs (middle) and microtubule that are color coded according to the local nearest distance to a neighboring microtubule, with red being 25nm and white larger than 100nm. Scale bar is 1µm. (Control 1 and 2 were previously shown in a different context ^7^ and have been adapted) **B.** Table of all quantified parameters for each dataset.

## Movies

**Movie 1. Related to Figure 2**

Time-lapse movie of a *C. elegans* one-cell embryo expressing EBP2 GFP and cherry Histone. Interval 2 sec.

**Movie 2. Related to Figure 2**

Time-lapse movie of a *C. elegans* one-cell embryo expressing EBP2 GFP and cherry Histone after *spd-1 (RNAi).* Interval 2 sec.

**Movie 3. Related to Figure 2**

Time-lapse movie of a *C. elegans* one-cell embryo expressing EBP2 GFP and cherry Histone after *spd-/ gpr-1/2 (RNAi).* Interval 2 sec.

**Movie 4. Related to Figure 2**

Time-lapse movie of a *C. elegans* one-cell embryo expressing EBP2 GFP and cherry Histone after *klp-19 (RNAi).* Interval 2 sec.

**Movie 5. Related to Figure 2**

Time-lapse movie of a *C. elegans* one-cell embryo expressing EBP2 GFP and cherry Histone after spd-1/ *klp-19 (RNAi).* Interval 2 sec.

**Movie 6. Related to Figure 4**

Time-lapse movie of metaphase cell-derived KLP-19::GFP moving on pre-polymerized, fluorescently labeled microtubules. Interval 1 sec. Still from this movie are shown in Figure 4B, Example 1

**Movie 7. Related to Figure 4**

Time-lapse movie of anaphase cell-derived KLP-19::GFP moving on pre-polymerized, fluorescently labeled microtubules. Interval 1 sec. Still from this movie are shown in Figure 4B, Example 2.

**Movie 8. Related to Figure 4**

Time-lapse movie of anaphase cell-derived SPD-1::GFP binding and concentrating on region of dense and brighter tubulin signal. Interval 1 sec. Still from this movie are shown in Figure 4E

## Acknowledgments

The authors are thankful to the members of the Molecular Electron Microscopy core at the University of Virginia for technical assistance, Y. Chen (UVA, Charlottesville) for initial help in *C. elegans* crossings.

M. Srayko (University of Alberta) for the MAS91 *C. elegans* strain, M. Delattre (University of Lyon) for the ANA72 strain and K. O’Connell for the *zyg-1* feeding clone. V.Z. and M.M. designed the experiments and conducted most of the light-microscopy experiments of RNAi and control conditions and analyzed the data. N.I.M. and D.J.D. designed, conducted and quantified all *in* vitro single cell lysate experiments, C.Y., M.B., D.N. and V.Z. designed, conducted and quantified the SHG experiments, X.H. conducted some light-microscopy and analysis of RNAi conditions, K.S. designed the mitotracker software to quantify the light microscopy data, A.P. and T.G. helped with the design of the KLP-19 GFP and SPD-1 Halo Tag strain and the injections as well as screenings, S.R. oversaw the experiments and acquired and quantified the tomographic data, V.Z. and S.R. co-wrote the paper, all other authors provided additional input and suggestions.

S.R. was supported by National Institutes of Health Grant (NIH) R01GM144668, X.H. was supported by NIH RGM144668A, M. M. was supported by the Wagner Fellowship, N.I.M. was supported by NIH F31 HD108006, C. Y. and D. N. were supported by Human Frontier Science Program (RGP 7 0034/2010), National Science Foundation (NSF) grant DMR-0820484 and NIH 1R01GM104976-01, T. G. was supported by NIH F31 HD112152, A. P. was supported by NIH R35 GM142880 and D. J. D. was supported by NIH R01 GM138443 and NIH R21 GM144817. The authors declare that they have no competing interests. All data needed to evaluate the conclusions in the paper are present in the paper and/or the Supplementary Materials. The light- and electron microscopic data can be provided by S.R. pending scientific review and a completed material transfer agreement. Requests for the data should be submitted to the corrisponding authors.

